# Evolutionary history, ecological divergence, and introgression in the *Oncocyclus* irises species complex in the Southern Levant

**DOI:** 10.1101/2025.10.16.682823

**Authors:** Yamit Bar-Lev, Sissi Lozada-Gobilard, Aleena Xavier, Lior Glick, Itay Mayrose, Yuval Sapir

**Affiliations:** The Botanical Garden, School of Plant Sciences and Food Security, G.S. Wise Faculty of Life Science, Tel Aviv University, Israel; Department of Biology, Lund University, Lund, Sweden; Tropical Ecology and Evolution Lab, Department of Biological Sciences, Indian Institute of Science, Education and Research, Bhopal, Madhya Pradesh, India; School of Plant Sciences and Food Security, G.S. Wise Faculty of Life Science, Tel Aviv University, Israel

**Keywords:** Phylogenetics, Royal Irises, *Oncocyclus*, IBD, IBE

## Abstract

Speciation is a dynamic process shaped by the interaction of gene flow, geographic isolation, and ecological divergence. The *Oncocyclus* irises of the Southern Levant represent a young radiation of narrowly endemic species considered to be in the course of speciation. In this study, we used RAD-sequencing and single nucleotide polymorphism analysis across nine described species to investigate patterns of genomic divergence, gene flow, and local adaptation. Phylogenomic analyses revealed a mix of well-supported clades for some species, previously defined by their morphology and distribution, and non-monophyletic lineages, with several species exhibiting shallow divergence and shared genetic ancestry. We found evidence for gene flow and historical introgression between *Iris petrana*, *I. atrofusca* and *I. mariae*, while other cases of non-monophyly appear driven by incomplete lineage sorting. Both geographic distance (IBD) and environmental factors (IBE), mainly altitude, temperature, and aridity, were significantly associated with genetic structure, suggesting that local adaptation contributed to divergence following range expansion. Based on our findings we propose that the divergence of the *Oncocyclus* iris species in the Southern Levant supports a stepping-stone dispersion model, in which north-to-south dispersal was followed by local adaptation, and introgression in secondary contact zones.

Overall, these findings highlight the complexity of speciation and the need for integrative approaches to study the interplay between historical divergence, contemporary gene flow, and environmental differentiation in shaping genomic patterns.

## Introduction

Speciation, the process by which populations diverge into distinct species through the accumulation of reproductive barriers, is of essential importance to evolutionary biology (Mayr 1942, Coyne and Orr 2004, Nosil 2012, Baack et al. 2015). Speciation is now recognized as a continuous process, with species existing at various stages of divergence (Nosil 2012, Stankowski and Ravinet 2021). This continuum is characterized by heterogeneous genomic divergence, where selection and gene flow interact to shape patterns of differentiation across the genome (Nosil 2012, Seehausen et al. 2014, Wolf and Ellegren 2017). Studying lineages at early stages of divergence, in which reproductive isolation is incomplete, can shed light on the mechanisms driving speciation.

Two major processes contribute to divergence: reduced gene flow and local adaptation (Thorpe et al. 2008, Nosil 2012). Reduced gene flow between geographically isolated populations leads to Isolation by distance (IBD), where genetic differentiation increases with geographic distance (Slatkin 1993, Moyle 2006, Jenkins et al. 2010, Sapir and Mazzucco 2012, Orsini et al. 2013, Guzmán et al. 2022). Alternatively, isolation by environment (IBE) occurs when ecological divergence reduces gene flow due to selection against maladapted migrants or hybrids (Shafer and Wolf 2013, Wang and Bradburd 2014). In this scenario, genetic divergence correlates with ecological differences (Rundle and Whitlock 2001, Orsini et al. 2013, Wang and Bradburd 2014). Notably, these processes often coexist, resulting in complex patterns of divergence (Feder et al. 2012, Guzmán et al. 2022).

Modern speciation research shows that genomic divergence is often heterogeneous across the genome, with some regions diverging due to selection or reduced recombination, while others remain similar due to ongoing gene flow (Seehausen et al. 2014, Wolf and Ellegren 2017, Stankowski and Ravinet 2021). This pattern helps identify genomic regions involved in adaptation and reproductive isolation, and reveals the dual role of gene flow in shaping divergence (Feder et al. 2012, Nosil 2012, Stankowski and Ravinet 2021). Gene flow can both counteract divergence by homogenizing populations, or promote it by facilitating adaptation and through introgression (Eaton et al. 2015, Tigano and Friesen 2016, Stankowski and Ravinet 2021, Bock et al. 2023). Hybridization and introgression are recognized as creative forces in diversification, generating novel traits and blurring species boundaries, resulting in complex patterns of genetic variation (Anderson and Stebbins 1954, Abbott et al. 2013, Harrison and Larson 2014, Bock et al. 2023). Species complexes, the groups of closely related taxa at varying divergence stages, are ideal systems for studying these dynamics (Struck et al. 2018, Osmolovsky et al. 2022). By analyzing genome-wide patterns of divergence and introgression in these complexes (e.g., using RAD-seq and ABBA-BABA tests), we can disentangle the roles of geography, ecology, and genomic architecture in shaping speciation (Eaton and Ree 2013, Martin et al. 2015, Campbell et al. 2018, Hibbins and Hahn 2022).

The Royal Irises (*Iris* section *Oncocyclus*, Baker) represent an ideal system for studying speciation, comprising approximately 30 species that are hypothesized to have diverged recently and are currently in various stages of speciation (Avishai and Zohary 1980, Sapir and Shmida 2002, Abdel Samad et al. 2016, Wilson et al. 2016). This species complex may have originated in the Caucasus mountains and spread throughout the eastern Mediterranean and the Levant (Wilson et al. 2016). It is especially valuable for speciation research because it exhibits multiple reproductive barriers. Intraspecific post-zygotic barriers have been documented, for example in *Iris atropurpurea* (Yardeni et al. 2016), while interspecific isolation is primarily pre-zygotic and ecogeographical, with weak or absent post-zygotic barriers (Volis et al. 2021, Osmolovsky et al. 2022).

These patterns point to a complex interplay of geographical, ecological, and genetic factors driving diversification within the group. Contrasting hypotheses have been proposed regarding the relative contributions of geographic isolation and ecological divergence in the speciation process of this young clade (Wilson et al. 2016, Yardeni et al. 2016). Morphological and genetic variation suggest that both factors may be involved (Sapir and Shmida 2002, Sapir and Mazzucco 2012, Yardeni et al. 2016). For instance, morphological gradients, such as a latitudinal cline in plant size, indicate adaptation to environmental conditions (Arafeh et al. 2002, Sapir et al. 2002). Recent studies suggest that ecogeographical reproductive barriers drive the speciation of the Royal Irises (Volis et al. 2021, Osmolovsky et al. 2022). Genetic studies in *Oncocyclus* suggested restricted gene flow among some populations but could not resolve intraspecific divergence or introgression (Arafeh et al. 2002, Saad et al. 2009, Wilson et al. 2016). However, these studies have not incorporated genome-wide genetic analyses or intra-specific sampling at sufficient resolution. Thus, the genomic patterns and processes underlying speciation in this group, and the relative contributions of geography and ecology, remain poorly understood.

Our study uses restriction-site associated DNA sequencing (RAD-seq) to retrieve genome-wide polymorphic sites across multiple populations and species of the *Oncocyclus* irises in Israel. This approach allows us to ask: (1) What are the phylogenetic and population genomic relationships within the *Oncocyclus* clade? (2) To what extent do current taxonomic species boundaries reflect underlying genomic structure? (3) Are patterns of genomic divergence shaped primarily by geography, ecological divergence, or gene flow? (4) Has historical introgression contributed to diversification or ecological differentiation within the group? By integrating phylogenetics, population structure analyses, and tests for IBD/IBE and introgression, we provide a genomic perspective on speciation mechanisms in this radiation.

## Methods

### Plant materials

For population sampling of the *Oncocyclus*, we used the current accepted taxonomy, following Flora of Israel and Adjacent Areas (https://flora.org.il/en/en/). We collected leaves from populations that represent the species’ spatial distribution (Figure 1). Samples of 68 *Iris* populations in Israel, Palestine, Jordan and Syria were included in the study, one sample per population (see map in Figure 1 and Suppl. Material Table S1). We also included two non-Oncocyclus *Iris* species to serve as outgroups: four samples from two populations of *I. lutescens* (obtained from Dr. E. Imbert, University of Montpellier, France) and one sample from a natural population of *I. mesopotamica* (collected in Mount Hermon, Israel). An overview of the species and populations represented in the sampling, as well as geographical and environmental characteristics of the populations, are detailed in Supplementary material Table S1.

**Figure 1.**
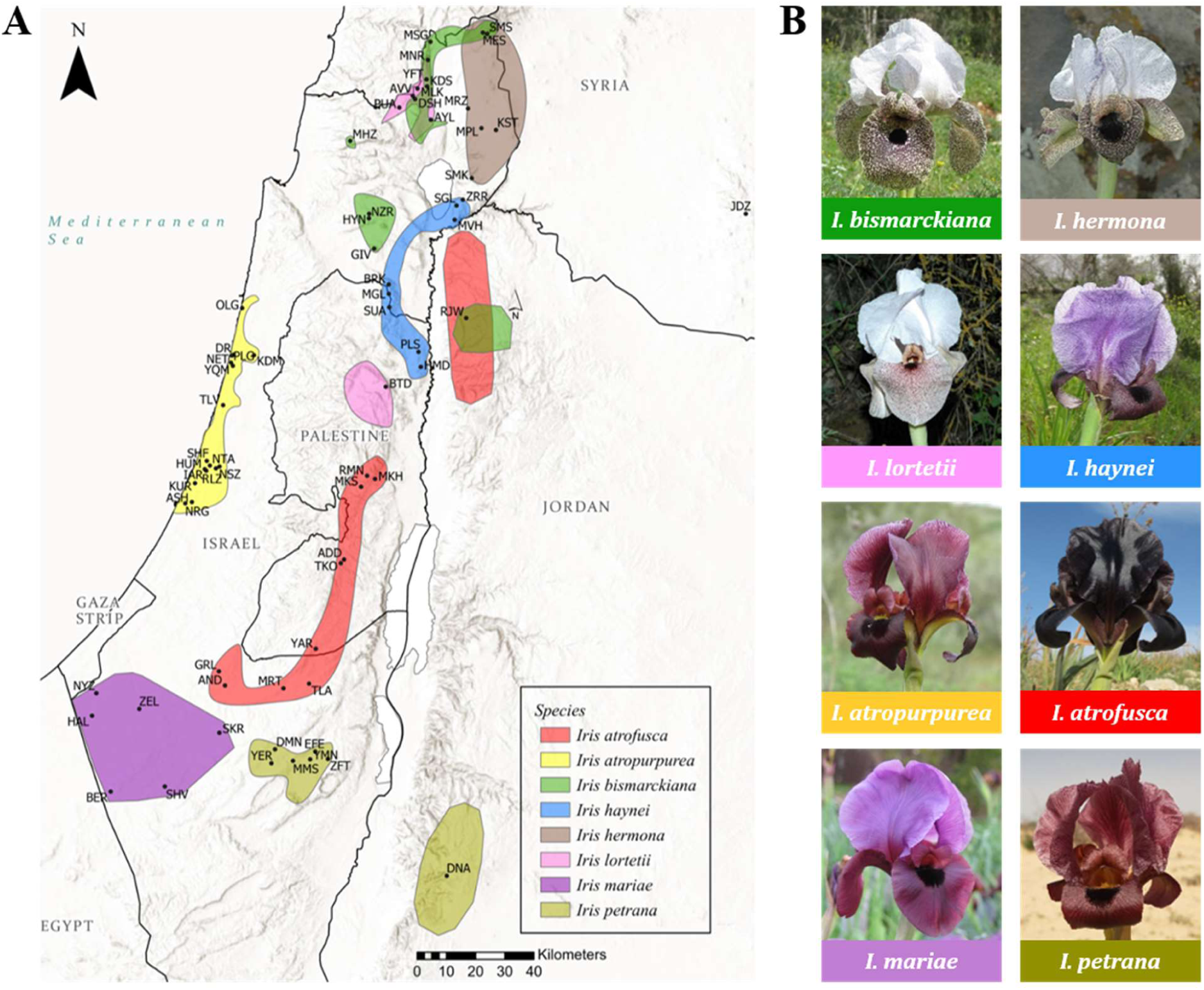
Populations sampled in the Southern Levant (Israel, Palestine, Jordan and Syria). **(A)** Coloured regions correspond to the natural distribution of the species (Y. Sapir, unpublished data) and the black dots correspond to the sampling locations. **(B)** Representative pictures of the Israeli *Iris* species. Pictures credits: Yuval Sapir.

### DNA extraction and RAD sequencing

We extracted genomic DNA from *Iris* leaves using CTAB protocol (Doyle and Doyle 1987) and DNeasy plant mini kit (Qiagen), following the manufacturer’s instructions. Library preparation and RAD sequencing were performed by Floragenex Inc. (Eugene, Oregon) using the restriction enzyme *PstI* and sample-specific barcodes. Samples were pooled and run multiplexed on two lanes of 1X100bp Illumina HiSeq 2000 sequencer.

### SNP calling

Floragenex Inc. processed the raw sequence data (Illumina FASTQ output files) using an internal pipeline. Assembly to contigs was performed using Velvet 1.2.10 (Zerbino and Birney 2008). The genome of *I. atropurpurea* from the Rishon LeZion population (RLZ) was *de novo* assembled into contigs due to its large number of high-quality reads (17,930,977 reads, 99.4%). The initial assembly of the reference individual was realigned to itself using BWA 0.6.1 (Li and Durbin 2009). Sequences were then aligned to the reference sequence using Bowtie 1.1.1 (Langmead et al. 2009). Variant calling was performed by mapping RAD-seq reads to the RLZ contigs using SAMtools 0.1.16 (Li 2011) and custom Floragenex scripts (average of 36.4% of reads aligned). We filtered the SNPs using the software SNPhylo (Lee et al. 2014) version 20140701. Specifically, we discarded monomorphic and multiallelic SNPs, as well as those with minor allele frequency below 0.01 or with missing data ratio above 0.5, and filtered the SNPs to avoid linkage disequilibrium.

The VCF file containing the final SNP data can be found in the Supplementary.

### Genetic structure

We conducted STRUCTURE analysis (Pritchard et al. 2000), using Bayesian approach to infer the number of genetic clusters (K). We tested K values from 2 to 10, with ten independent runs for each K. Each run included a burn-in of 200,000 iterations and a total of 200,000 Markov Chain Monte Carlo iterations. We determined the most probable K using ΔK values, calculated in Structure Harvester (Evanno et al. 2005, Earl and VonHoldt 2012). We used CLUMPAK to summarize and visualize results (Kopelman et al. 2015). Unlike standard STRUCTURE input, we treated populations as “individuals” and species as “populations”, to reflect species-level structure.

We also performed a discriminant analysis of principal components (DAPC) using the R package adegenet (Jombart 2008). We retained 30 principal components in the DAPC, based on the cumulative variance plot, to balance information retention and avoid overfitting. Discriminant analysis was then performed using the first 4 linear discriminants. In both STRUCTURE and DAPC analyses, *I. bismarckiana* was genetically distinct, skewing the results. To enable fine scale resolution of the remaining populations, both analyses were performed also excluding outgroups and *I. bismarckiana* populations (n=59).

### Phylogenetic analysis

A species tree was inferred based on the SNP data using the software SNPhylo (Lee et al. 2014) version 20140701 with parameters “-A -b -p 90 -M 0.5 -m 0.01”. A maximum likelihood tree was inferred using DNAML from the PHYLIP R package (Revell and Chamberlain 2014). Bootstrap analysis was performed using the Phangorn R package (Schliep 2011). Although two outgroup species were included in the dataset, the phylogenetic tree was rooted using only *I. mesopotamica* because it is more closely related to the ingroup taxa than *I. lutescens*, minimizing long-branch attraction and better reflecting the likely root position (Abdel Samad et al. 2016). The tree was visualized using FigTree v1.4.3 (Rambaut 2007).

To quantify genetic differentiation among species, we also calculated pairwise F_ST_ values using the Weir and Cockerham (1984) estimator (Weir and Cockerham 1984). Calculations were performed in R using the genet.dist() function from the hierfstat package, applied to a filtered genind object containing individuals grouped by species. The resulting F_ST_ matrix was rounded to three decimal places and visualized as a heatmap to highlight levels of genomic differentiation between taxa.

### Tests for introgressions

We tested for introgressions among taxa using Patterson’s D statistics (Durand et al. 2011), also called the ABBA-BABA test, in D-suite (Malinsky et al. 2021). This test uses three taxa (P1, P2, P3) and a putative outgroup (O) and quantifies the number of shared and derived alleles, as well as tests for introgression between the three taxa, by calculating the difference between frequencies of two allele patterns across all SNPs shared by the four groups. Denoting A as the ancestral allele and B as the derived allele, their order in the tree can be ABBA or BABA:

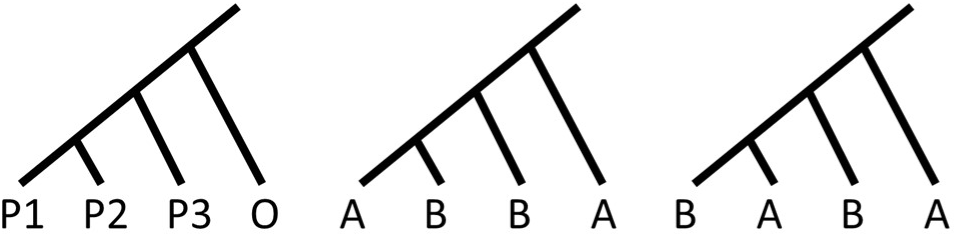

Under random incomplete lineage sorting (ILS) scenario, both ABBA and BABA patterns are equally likely for each locus. The overrepresentation of ABBA pattern indicates gene flow between P3 and P2, whereas the overrepresentation of BABA pattern indicates gene flow between P3 and P1.

We have constructed a few hypotheses regarding introgressions among taxa, based on incongruences between the genetic clustering pattern in the reconstructed phylogenetic tree (Figure 3) and either the geographical distribution or the recognized taxonomy, which was derived based on morphological data (Sapir et al. 2002, Danin and Fragman-Sapir 2016+). In most tests, we used *I. bismarckiana* as an outgroup, and when the hypothetical introgression involved *I. bismarckiana*, we used *I. mesopotamica* as an outgroup (Table 1).

**Table 1.** List of the ABBA-BABA tests performed based on the SNP data. D-stat: Patterson’s D statistics, the normalized differences between ABBA and BABA patterns. Z-score: how many standard differences D is from zero. F4 ratio: admixture proportion, BBAA: frequency of alleles showing BBAA pattern, ABBA: frequency of alleles showing ABBA pattern, BABA: frequency of alleles showing BABA pattern.

| No. | P1 | P2 | P3 | O | D-stat | Z-score | p-value | f4-ratio | BBAA | ABBA | BABA |
| --- | --- | --- | --- | --- | --- | --- | --- | --- | --- | --- | --- |
| 1 | *I. haynei* s.s. | *I. haynei* s.l. | PLS + HMD (*I. haynei*) | *I. atrofusca* | 0.026 | 1.625 | 0.104 | 0.016 | 304.15 | 200.87 | 190.61 |
| 2 | *I. lortetii* | *I. haynei* s.s. | RMN+MKS+MKH<br>(*I. atrofusca*) | *I. bismarckiana* | 0.017 | 0.786 | 0.432 | 0.014 | 278.38 | 251.25 | 243.01 |
| 3 | *I. hermona*<br>(excl. SMK) | SMK | *I. bismarckiana* | *I. mesopotamica* | 0.083 | 1.471 | 0.141 | 0.037 | 72.27 | 61.96 | 52.50 |
| 4 | *I. atrofusca* | *I. hermona* | *I. bismarckiana* | *I. mesopotamica* | 0.114 | 2.755 | **0.006** | 0.041 | 81.00 | 58.28 | 46.39 |
| 5 | *I. atropurpurea* | *I. mariae* | *I. atrofusca* | *I. bismarckiana* | 0.064 | 4.990 | **<0.0001** | 0.128 | 282.29 | 259.40 | 228.07 |
| 6 | *I. atropurpurea* | *I. mariae* | *I. petrana*+ *I. atrofusca*+<br>PLS + HMD (*I. haynei*) | *I. bismarckiana* | 0.080 | 6.101 | **<0.0001** | 0.176 | 279.98 | 265.61 | 226.26 |
| 7 | *I. atrofusca* | *I. petrana* | *I. mariae* | *I. bismarckiana* | 0.061 | 3.845 | **<0.0001** | 0.061 | 330.69 | 251.88 | 223.05 |

### Genetic, geographical and ecological distances

We built three matrices based on genetic, geographical and environmental distances between all populations. The genetic matrix was based on the SNP dataset after filtering (total number of SNPs=28,713), and calculated using the cophenetic distances of the hierarchical clustering. The geographical matrix was based on the spatial coordinates (longitude and latitude) using Euclidean distances. The environmental matrix was built considering the following variables: altitude, mean annual temperature, mean annual precipitation, annual evaporation, the minimal temperature in February, aridity and soil type (Suppl. Material Table S1). For soil type, we ran a Principal Component Analysis (PCA) with 10 binary variables of soil types, and the first component, explaining 61% of the variance, was included as a soilPC1 variable. Highly correlated predictors (|r|>0.75) were removed (Suppl. Material Figure S3). After the selection, five variables were included: altitude, mean annual temperature, annual evaporation, aridity and SoilPC1. We calculated all distances for the three matrices using the function *dist* from the R package “stats” with the *euclidean* method.

### Isolation by distance and isolation by ecology

We performed IBD and IBE analyses only for the *Iris* species growing in Israel, excluding the outgroup species, *I. lutescens* and *I. mesopotamica*. We also excluded *I. auranitica* (JDZ), the Jordanian populations of *I. petrana* (DNA) and *I. bismarckiana* (RJW), and two more populations (AVV and SMS) due to a lack of environmental data (Suppl. Material Table S1). In addition, one species, *I. bismarckiana*, formed a clear separated group in the STRUCTURE, DAPC, and the phylogeny (see Figures 2A, 3, and Suppl. Material Figures S1A and S2), which may bias the results. Thus, all tests for IBD and IBE were performed twice, once with all species (n=63) and once excluding *I. bismarckiana* (n=56). Isolation by distance (IBD) was evaluated by testing the correlation between genetic and geographical matrices using Mantel tests. The Mantel test is a statistical method to examine the relationship between geographic distance and ecological similarity (Mantel 1967). Similarly, Isolation by Environment (IBE) was evaluated by comparing the genetic matrix with the environmental matrix. Correlation significance tests were carried out using the mantel.randtest function in the R package *adegenet* (Jombart 2008), with 10,000 permutations.

**Figure 2.**
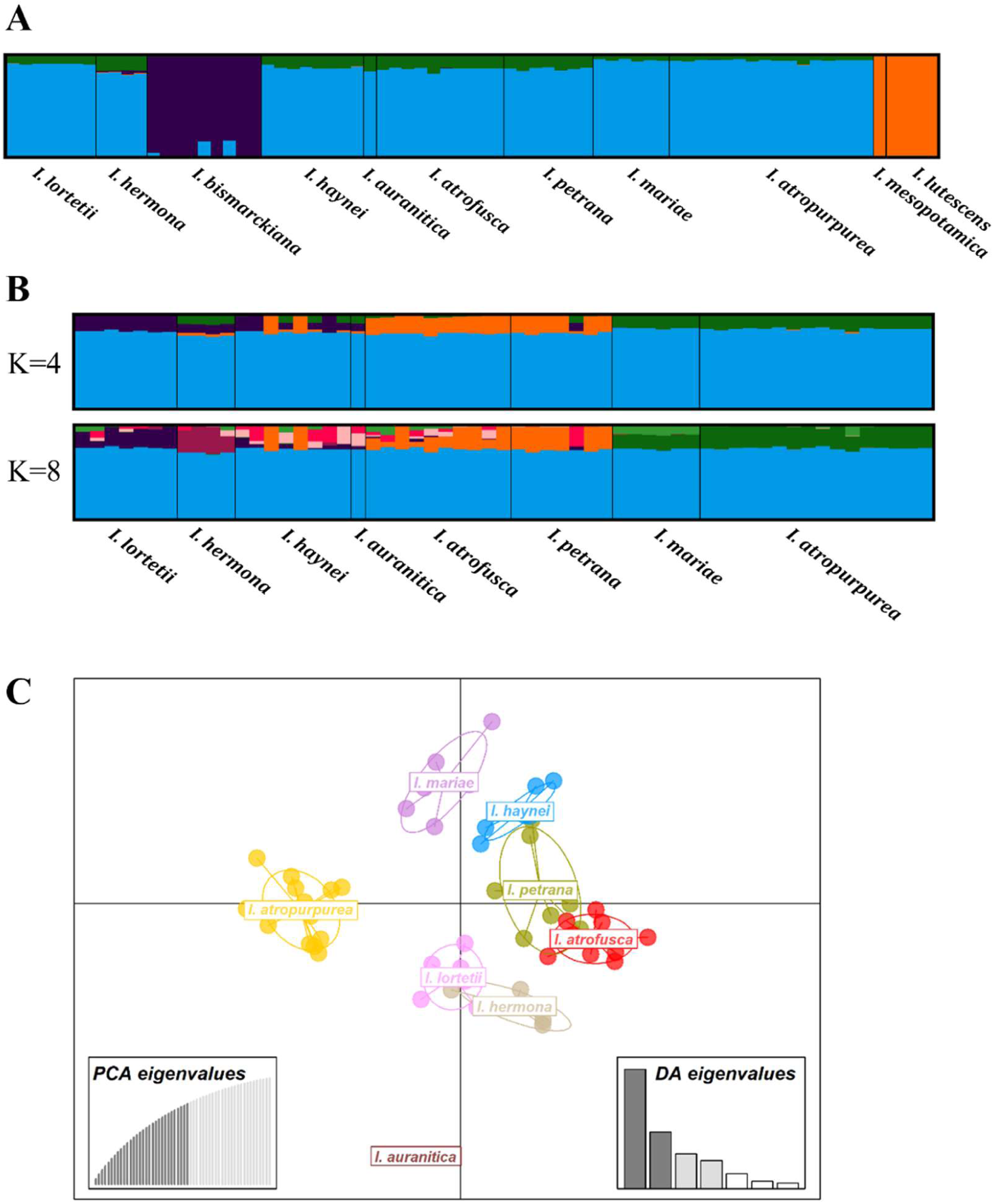
Genetic structure of *Iris* species based on STRUCTURE and DAPC analyses. **(A)** STRUCTURE plot at K = 4 including outgroups (*I. lutescens*, *I. mesopotamica*) and *I. bismarckiana* (n=73), showing distinct clusters for these species and a shared ancestry among the remaining taxa. **(B)** STRUCTURE plots at K = 4 and K = 8 excluding outgroups and *I. bismarckiana* (n=59), revealing shared ancestry with finer-scale structure and species-specific components. **(C)** Discriminant Analysis of Principal Components (DAPC) excluding *I. bismarckiana*, showing clear clustering among most species with some overlap between *I. petrana* and *I. atrofusca*, and between *I. hermona* and *I. lortetii*.

Due to some correlation between the geographical and ecological distances, we also performed partial Mantel test to examine the relative effect of geographical and environmental distances on the genetic distance. In this test, the correlation between two matrices is considered while accounting for the third one, i.e., the correlation between the genetic and geographical distances accounting for the environment, as well as the correlation between the genetic and environment distances, accounting for the geographical distance. Partial Mantel tests were performed with the function mantel.partial from the R package *vegan* (Oksanen et al. 2013).

Redundancy analysis (RDA) was performed to understand which environmental factors best explain the genetic variation (reviewed in (Capblancq and Forester 2021)). RDA is an ordination, based on multivariate regression that identifies linear association of the explanatory variables (i.e., environmental predictors) that maximizes the explained variance of the response variables (i.e., SNPs). Models were constructed with the five uncorrelated environmental predictors (see above) and tested for multicollinearity using Variance Inflation Factors (VIF), where values lower than 10 are used for further analyses (Dyer et al. 2010, Zuur et al. 2010). To test which environmental variables are significantly associated with the SNPs, we applied the function “anova.cca” with the option by “terms” from the *vegan* package.

## Results

### RAD-sequencing and SNP calling

To evaluate species structure and phylogeny, we RAD sequenced representative samples of the *Iris* species, which generated 586,253,421 reads. *De novo* assembly resulted in a total of 98,918 assembled contigs, each 92 bp in length, with total assembly length of ∼9.1 Mb. The number of RAD clusters for each species is elaborated in Supplementary material Table S1. Variant calling resulted in 310,796 candidate polymorphic loci, out of which 28,713 were retained after filtering.

### Population genetic structure

Population structure analysis using STRUCTURE and Discriminant Analysis of Principal Components (DAPC) revealed both shared ancestry and species-specific differentiation among *Iris* species. STRUCTURE analysis inferred four ancestral genetic clusters, with *I. lutescens* and *I. mesopotamica*, and *I. bismarckiana* forming distinct groups, while all other species shared a common genetic background (Figure 2A; the full STRUCTURE plot is in the Suppl. Material Figure S1A). After excluding the three taxa, the optimal number of ancestral groups remained four, with a predominant cluster (light blue) consistently present in all species, indicating shared ancestry. Distinct genetic components were shared between *I. atropurpurea* and *I. mariae* (green), showing homogeneous profiles with little admixture, as well as between *I. atrofusca* and *I. petrana* (orange). One distinct component (purple) was most prominent in *I. lortetii*, while *I. haynei* exhibited the most intra-species variability (Figure 2B; the full STRUCTURE plot is in the Suppl. Material Figure S1B). At k=8, which is the number of taxonomically described species in Israel, a similar pattern was observed, with additional fine-scale population structure. Most notably, unique genetic component became prominent in *I. mariae* and *I. atropurpurea* (light green), interrupting their previously homogeneous composition, and in *I. hermona* (burgundy). Additional components became visible across the other species, suggesting historical gene flow or complex divergence (Figure 2B).

DAPC also revealed *I. bismarckiana* as a highly distinct and separated cluster, indicating strong genetic differentiation. The remaining species formed adjacent clusters with partial overlap (Suppl. Materials Figure S2). To better resolve the relationships among the more closely related taxa, *I. bismarckiana* was excluded from the analyses. A total of 30 principal components were retained, explaining 75.7% of the total genetic variance, and the first four linear discriminants accounted for 89.2% of the among-group variation. The DAPC revealed well-defined and spatially distinct clusters for *I. atropurpurea*, *I. mariae*, and *I. auranitica*, suggesting high genetic coherence. Partial overlap was observed between *I. petrana* (olive-brown) and *I. atrofusca* (red), as well as between *I. hermona* (grey) and *I. lortetii* (pink), suggesting close genetic relationships and incomplete differentiation.

Together, these results highlight both deep and shallow genetic structure within Israeli *Iris* species, shaped by shared ancestry, recent divergence, and possible historical gene flow or hybridization.

### Phylogenetic analysis

To infer phylogenetic relationships among *Oncocyclus* irises, we reconstructed a maximum likelihood tree based on SNP data, rooted with *I. mesopotamica* (Figure 3). The outgroup species, *I. mesopotamica* and *I. lutescens*, formed a distinct clade, confirming their position outside the *Oncocyclus* group. Within the ingroup, *I. bismarckiana* emerged as a genetically distinct lineage, forming a well-supported clade separate from all other taxa. One population of *I. hermona* (SMK) was unexpectedly placed near *I. bismarckiana* and outside the main *I. hermona* clade, suggesting either misidentification or ancestral gene flow. *I. atropurpurea* and *I. mariae* each formed well-supported monophyletic clades, clearly distinct from other species. *I. lortetii* also formed a monophyletic clade, though with low bootstrap support. In contrast, *I. petrana*, *I. atrofusca,* and *I. haynei* showed non-monophyletic and intermixed relationships. *I. petrana* is polyphyletic: its Israeli populations nested within *I. atrofusca*, while the Jordanian population (DNA) grouped with *I. auranitica* from Syria (JDZ). *I. haynei* was also polyphyletic: three populations from Mt. Gilboa (SUA, BRK and MGL, hereafter *I. haynei* sensu stricto (s.s.)), formed a separated clade closer to *I. lortetii*, three other populations (MVH, SGL and ZRR, hereafter *I. haynei* sensu lato (s.l.)) clustered with the Jordanian and Syrian populations, and the other two populations (HMD and PLS) clustered with *I. petrana* and *atrofusca*.

**Figure 3.**
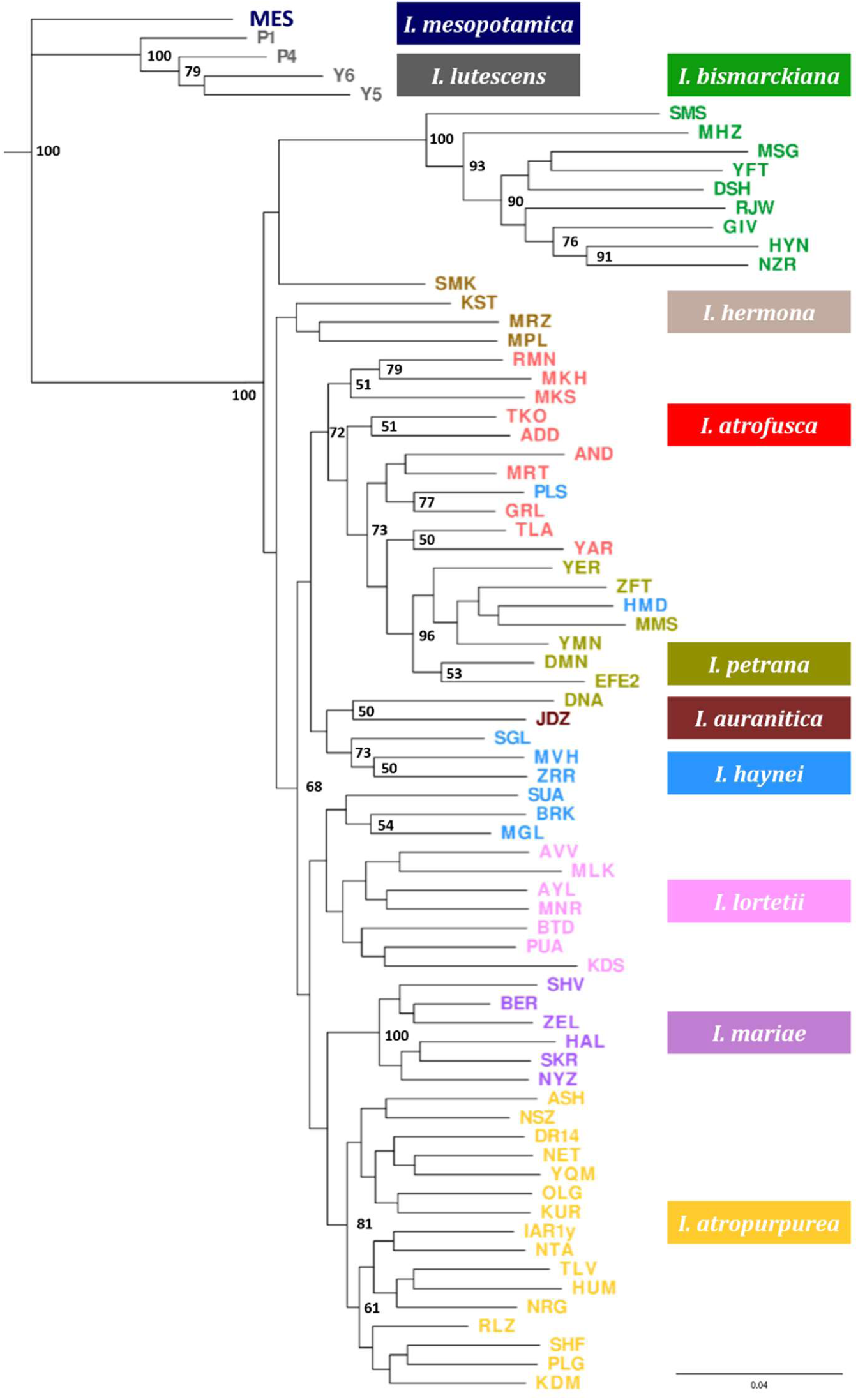
Phylogeny of the *Oncocyclus* irises. Maximum-likelihood phylogenetic tree inferred from the 28,713 SNPs. The tree is rooted with *Iris mesopotamica* and includes *I. lutescens* as an additional outgroup. Bootstrap values of >50 is indicated below the nodes.

To quantify genetic differentiation between species observed in the phylogenetic tree, we calculated pairwise F_ST_ values (Figure 4). The results corroborated the tree structure, with the highest F_ST_ values between *I. bismarckiana* and all other species (F_ST_ > 0.12). In contrast, very low F_ST_ values were found among *I. haynei*, *I. petrana*, and *I. atrofusca* (F_ST_ < 0.05), supporting the observed admixture and phylogenetic polyphyly within this group.

**Figure 4.**
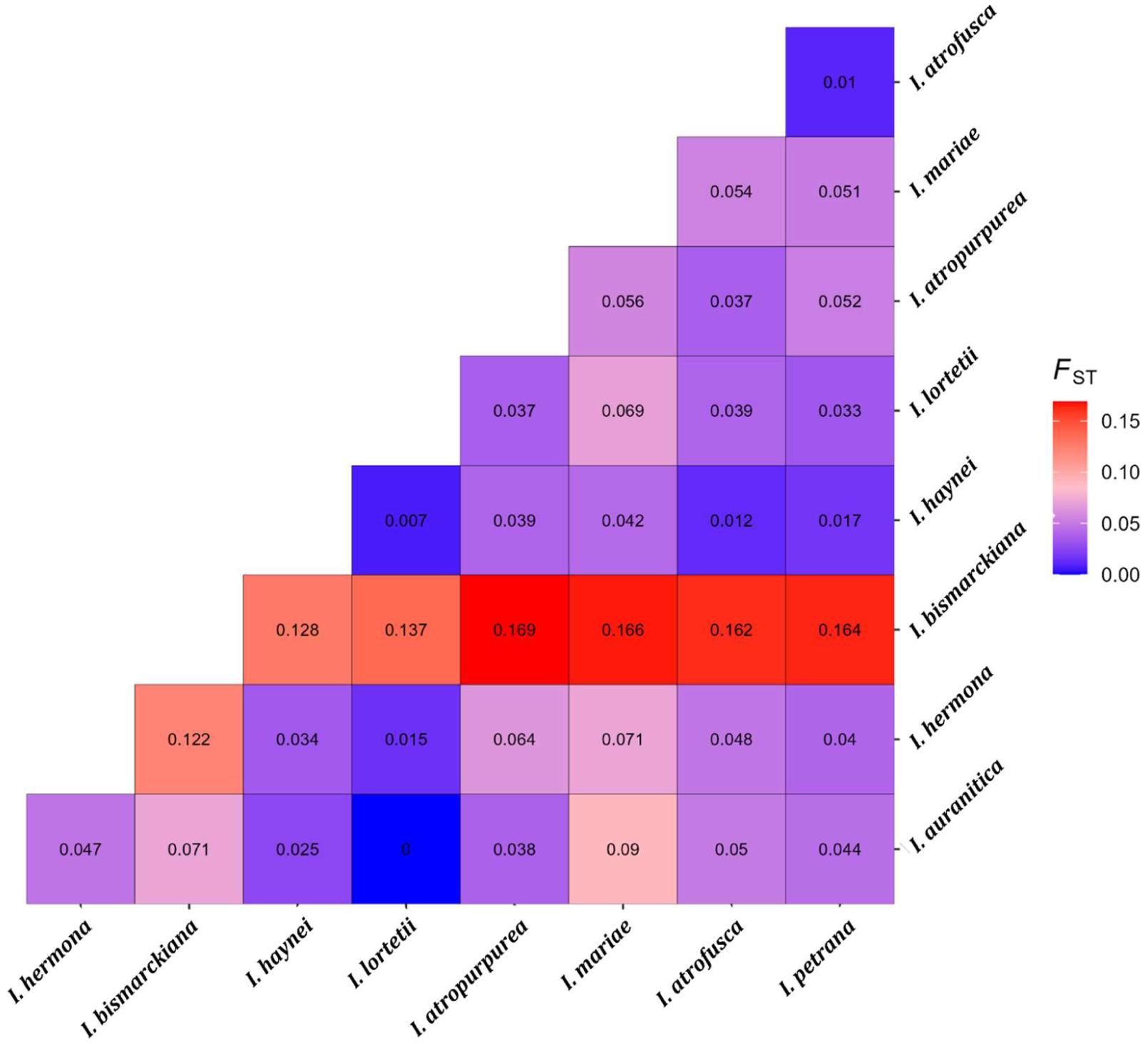
Pairwise F_ST_ values calculated using the Weir and Cockerham (1984) estimator, based on genome-wide SNP data. Warmer colors indicate higher genetic differentiation. Pairwise F_ST_ values ranged from 0.007 to 0.169, with the highest values observed between *I. bismarckiana* and other species.

These patterns indicate clear genetic structure for some taxa (e.g., *I. atropurpurea*, *I. mariae*, *I. bismarckiana*), while others show signatures of incomplete lineage sorting, introgression, or potential misclassification, particularly in the *I. petrana–atrofusca–haynei* complex.

### Tests for introgressions

To test for historical introgressions among *Iris* species, we conducted ABBA-BABA analyses (Table 1). Some tests, performed to investigate non-monophyletic structures observed in the phylogeny, did not show deviation from expectations under incomplete lineage sorting (ILS). For example, comparisons among *I. haynei* populations, which did not form a monophyletic group in the phylogeny, revealed no significant gene flow between southern populations (PLS, HMD) and either *I. haynei* s.s. (Gilboa populations) or the northern s.l. populations (Table 1, test 1). We also explored the unexpected phylogenetic clustering of *I. haynei* s.s. with *I. lortetii*, despite prior morphological evidence linking *I. haynei* to *I. atrofusca* (Arafeh et al. 2002, Sapir et al. 2002). We tested whether this grouping could result from introgression involving *I. atrofusca* populations but found no significant gene flow (Table 1, test 2). Together, these results support the interpretation that the relationships among *I. haynei* s.s., *I. lortetii*, and the southern *I. haynei* populations reflect deeper evolutionary history and lineage divergence, rather than recent gene flow.

To understand the phylogenetic placement of *I. hermona* population SMK, which clustered with *I. bismarckiana*, we tested for introgression of *I. bismarckiana* and *I. hermona* populations. There was no significant gene flow observed between *I. bismarckiana* and SMK directly (Table 1, test 3). However, significant excess allele sharing was observed between *I. bismarckiana* and *I. hermona* (D = 0.114, Z = 2.76, *p* = 0.006; Table 1, test 4), suggesting historical gene flow between these species. Notably, when *I. mesopotamica* was used as the outgroup (Tests 3-4), the number of shared loci dropped down (as low as 20%), likely due to its greater genetic divergence. Therefore, these results, recovered while using *I. mesopotamica* as the outgroup, should be interpreted with caution.

To assess the extent of introgression among southern desert taxa, we performed a series of ABBA-BABA tests involving *I. mariae*, *I. atrofusca*, and *I. petrana* (Table 1, tests 5–7). Significant allele sharing between *I. mariae* and *I. atrofusca* (D = 0.064, Z = 4.99, *p* < 0.0001; Table 1, test 5; Figure 5A) suggested direct gene flow or shared ancestry. When expanded to include *I. petrana* and southern *I. haynei* populations in a broader desert clade, the signal strengthened further (D = 0.080, Z = 6.10, *p* < 0.0001; Figure 5B), indicating more complex historical gene flow across species boundaries. Finally, Test 7 confirmed significant introgression between *I. mariae* and *I. petrana* (D = 0.061, Z = 3.85, *p* < 0.0001; Table 1, test 6; Figure 5C), suggesting that *I. mariae* has exchanged alleles with multiple taxa in the southern region, rather than with a single species.

**Figure 5.**
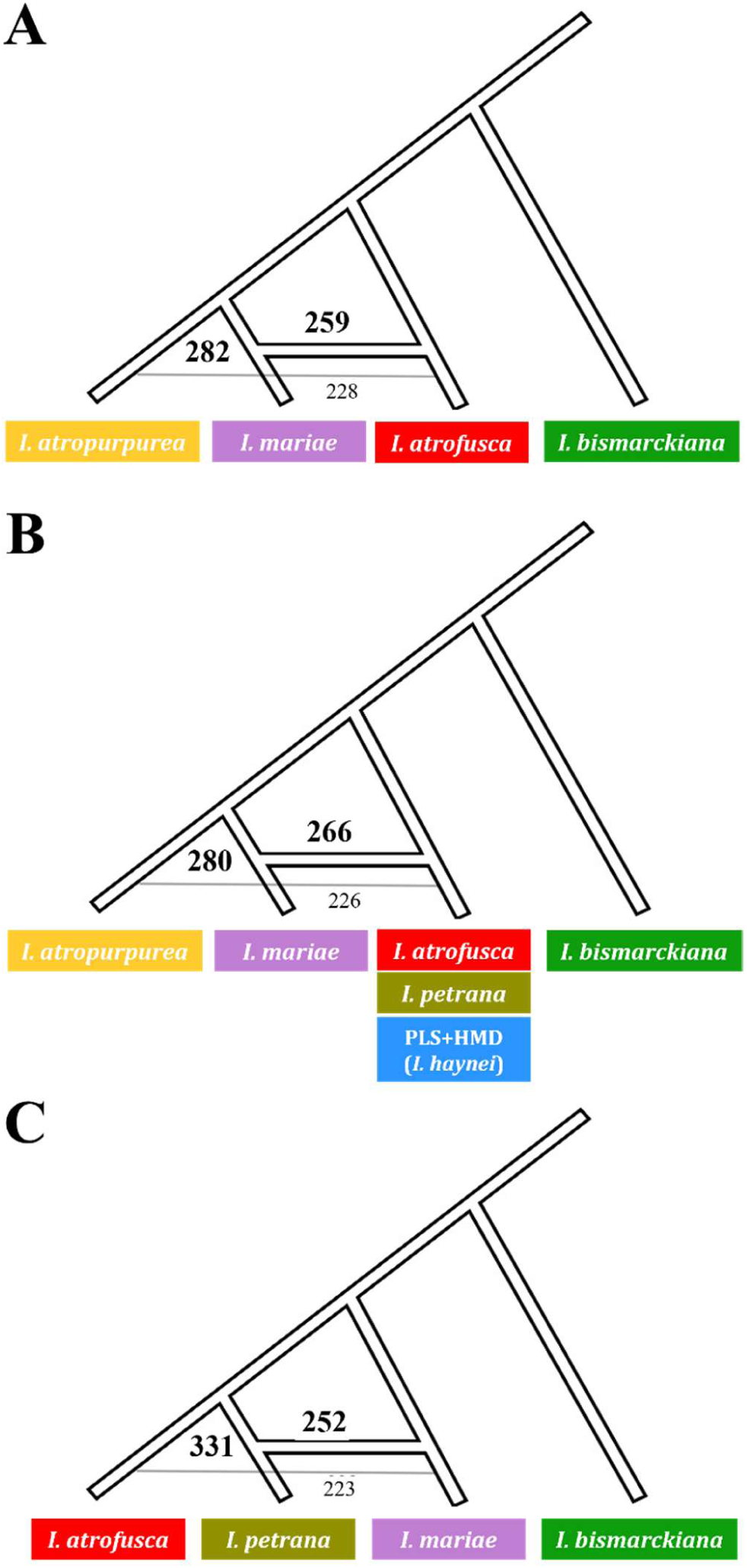
Introgression between **(A)** *Iris atrofusca* and *I. mariae* (BBAA-282, ABBA-259, BABA-228), **(B)** *I. atrofusca, I. petrana*, and *I. haynei* with *I. mariae* (BBAA-331, ABBA-252, BABA-223), and **(C)** between *I. mariae* and *I. petrana* (BBAA-280, ABBA-266, BABA-226). The three tests were performed with *I. bismarckiana* as an outgroup. Numbers indicate the frequencies of the different allele patterns.

### Isolation by distance and isolation by ecology

We tested for Isolation by distance (IBD) and Isolation by environment (IBE), and found a significant association between the genetic distances and both the environmental and geographical distances (p<0.05; Table 2; Suppl. Material Figure S4A-B). When we repeated the analysis excluding *I. bismarckiana* to improve resolution, the association was even more significant (p<0.001; Table 2; Figure 6A-B).

**Figure 6.**
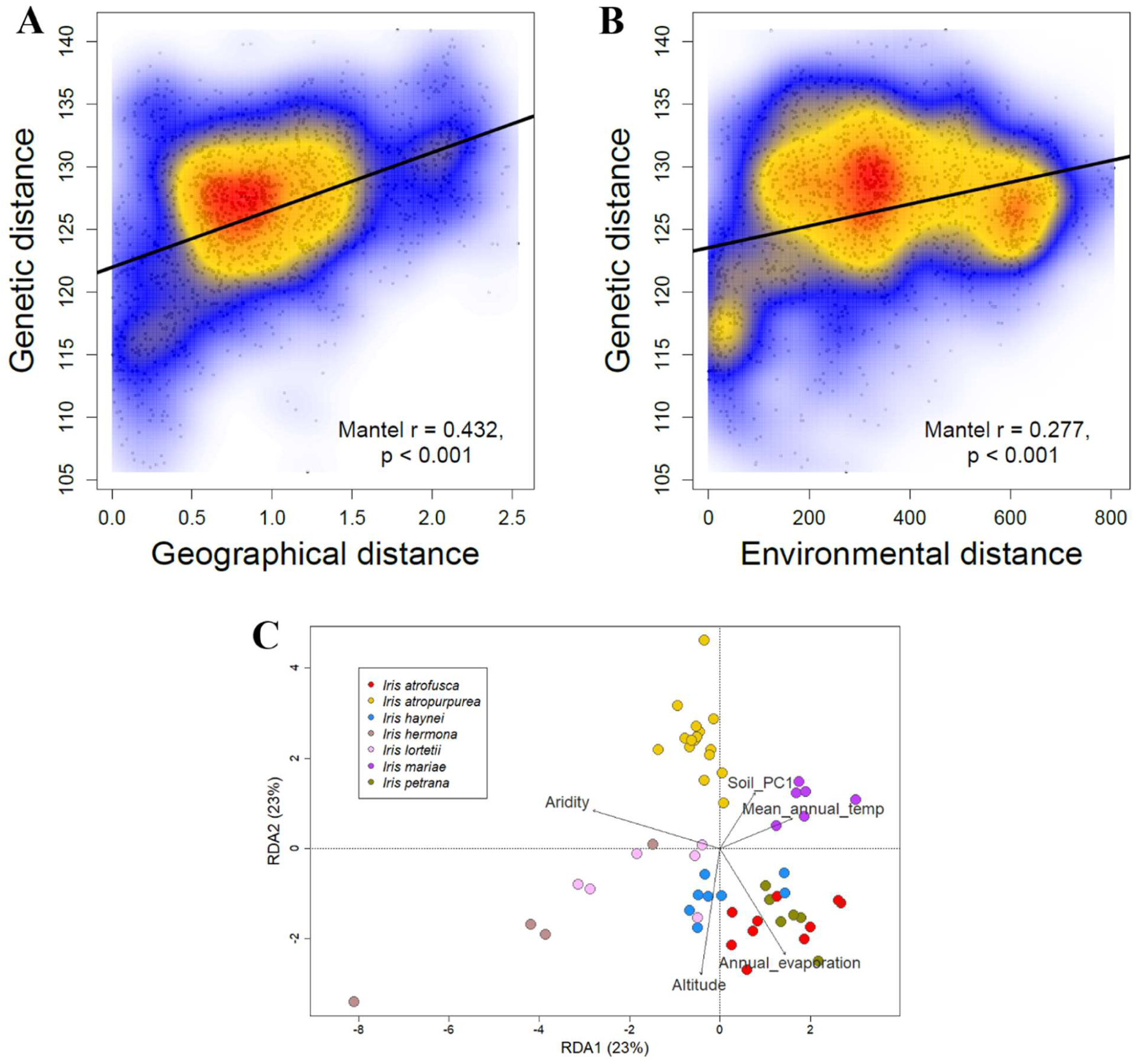
Isolation by distance **(A)** and Isolation by environment **(B)** were calculated by Mantel test excluding *I. bismarckiana* (n-56). Each dot represents a pair of populations, with position indicating their genetic distance and geographical or environmental similarity. Colors denote density of points. The solid line within each panel is the linear regression line. Mantel coefficient (ρ) of correlation between the matrices, and the p-values are displayed at the bottom right-hand side of each panel. **(C)** Redundancy analysis (RDA) ordination plot of the environmental variables based on genetic distance between the *Iris* species excluding *I. bismarckiana*. Each point represents a population, colours indicate the species, and the arrows represent the environmental variable. The length and direction of the arrows indicate the strength and direction of the relationships between the environmental variables and the genetic distance.

**Table 2.** Results of Mantel and partial Mantel tests for Isolation by distance (IBD) and Isolation by environment (IBE).

|  | Test | All species (n=63) |  | Excl. *I. bismarckiana* (n=56) |  |
| --- | --- | --- | --- | --- | --- |
|  |  | Mantel statistic *r* | *P*-value | Mantel statistic *r* | *P*-value |
| Mantel tests | IBD | 0.226 | <0.001 | 0.432 | <0.001 |
|  | IBE | 0.172 | 0.044 | 0.277 | <0.001 |
| Partial Mantel tests | IBD \| Environment | 0.192 | <0.001 | 0.397 | <0.001 |
|  | IBE \| Geographical | 0.123 | 0.099 | 0.210 | <0.001 |

To determine the individual contribution of geography and environment, we performed Partial Mantel tests. These showed a significant positive correlation between genetic and geographical distances when controlling for environmental variation, regardless of whether *I. bismarckiana* was included (p<0.001; Table 2). In contrast, the correlation between the genetic and environmental distances was only significant when *I. bismarckiana* was excluded (p<0.001; Table 2).

To test which environmental variables were most associated with the genetic distance, we performed redundancy analysis (RDA). There was no evidence for multicollinearity (all values < 10) in either dataset (with and excluding *I. bismarckiana*). The first two RDA axes explained 50% of the variation for all species (RDA1=28%, RDA2=22%; Table 3; Suppl. Material Figure S4C), and explained 46% when *I. bismarckiana* was excluded (RDA1=23%, RDA2=23%; Table 3; Figure 6C). In the reduced dataset, without *I. bismarckiana*, altitude, mean annual temperature and aridity were significantly associated with genetic distance (Table 3; Figure 6C).

**Table 3.** Effect of environmental predictors on the genetic relatedness of the Royal Irises complex. Each row indicates the explained genetic variation of the tested environmental variable, the associated Anova F statistic, and the p-value, as obtained with random permutations (*n*=9999) of the RDA model. Statistically significant results (α≤0.05) are in bold type.

|  | All species |  |  |  | Excl. *I. bismarckiana* |  |  |  |
| --- | --- | --- | --- | --- | --- | --- | --- | --- |
|  | Df | Variance | F | P | Df | Variance | F | P |
| Altitude | **1** | **476.6** | **1.28** | **0.001** | **1** | **476.7** | **1.31** | **0.001** |
| Mean annual temperature | 1 | 425.5 | 1.14 | 0.062 | **1** | **445** | **1.22** | **0.014** |
| Annual evaporation | **1** | **508.6** | **1.37** | **0.004** | 1 | 397 | 1.09 | 0.132 |
| Aridity | 1 | 419.6 | 1.13 | 0.104 | **1** | **424** | **1.17** | **0.049** |
| Soil type (PC1) | 1 | 349 | 0.94 | 0.798 | 1 | 358.4 | 0.98 | 0.546 |
| Residual | 57 | 21158.6 |  |  | 50 | 18114 |  |  |

## Discussion

Speciation is increasingly recognized as a dynamic and heterogeneous process, shaped by the combined effects of gene flow, ecological differentiation, and historical contingency (Nosil 2012, Seehausen et al. 2014, Stankowski and Ravinet 2021). In the early stages of divergence, genomic data often show a complex pattern, where some lineages are clearly distinct, while others remain mixed due to shared ancestry or gene flow (Feder et al. 2012, Seehausen et al. 2014, Wolf and Ellegren 2017). By examining a young radiation with varying levels of divergence and ecological differentiation, and by integrating genome-wide patterns of relatedness, this study illustrates how geographic distance, ecological gradients, and hybridization interact to shape evolutionary trajectories. We applied RAD sequencing across multiple *Oncocyclus* populations to resolve fine-scale genetic structure, reconstruct phylogenetic relationships, and identify signatures of demographic divergence and local adaptation.

### Genomic signatures of early divergence

In this study, we investigated the structure and genetic relatedness of populations belonging to nine recognized *Oncocyclus* irises species in the southern Levant. Our integrative population genomic analyses, combining STRUCTURE, DAPC, phylogenetic reconstruction, and F_ST_ estimates, revealed varying levels of genomic differentiation. We uncovered well-supported clades with clear species boundaries, such as *I. bismarckiana*, which appears genetically distinct across all analyses. *Iris mariae* and *I. atropurpurea* also show distinct genetic composition, characterized by their predominantly homogeneous genetic structure, indicating possibly stronger reproductive isolation. In contrast, other species complexes, such as *I. lortetii* and *I. hermona*, or *I. petrana* and *I. atrofusca*, show incomplete separation and substantial admixture patterns. This suggests historical and possibly ongoing gene flow, likely facilitated by their geographic proximity. In STRUCTURE analysis we found patterns a shared ancestry among species, further supporting recent common ancestry ILS, or historical gene flow.

Genomic divergence during speciation is often heterogeneous, with some regions of the genome showing high differentiation due to selection or reduced recombination, while others remain similar as a result of ongoing gene flow (Seehausen et al. 2014, Wolf and Ellegren 2017, Stankowski and Ravinet 2021). This pattern is expected in the “speciation continuum,” where populations are diverging but not yet fully isolated. This is further supported by the cytogenetic stability observed across *Oncocyclus* irises despite their extensive morphological divergence across the eastern Mediterranean, suggesting limited genomic change accompanying speciation (Abdel Samad et al. 2016).

Although Oncocyclus irises are cytogenetically stable (Abdel Samad et al. 2016), genome size evolution may still have played a role in their diversification. A recent study (Samad et al. 2020) found that genome size changes in Oncocyclus follow a speciation model, with shifts occurring primarily during speciation events. This finding aligns with our observation of distinct clades and genomic clusters, suggesting that subtle genomic shifts may accompany divergence even in the absence of large-scale chromosomal changes.

### The challenges of taxonomic resolution in young radiations

Most populations were segregated into clades that mostly align with the accepted taxonomy for the majority of the species (Figure 3). However, we did not observe a strong link between the phylogeny and the geographical distribution nor the species’ morphological delimitation. Moreover, although the accepted taxonomy consists of nine species, the STRUCTURE analysis indicates that the optimal number of genetic groups (K) is four. This suggests that some of the species are not genetically distinct, due to low divergence. Furthermore, the current taxonomy, which is based on morphological characteristics, may not accurately reflect the evolutionary history of the group, nor the true patterns of divergence and speciation. For example, despite the distinct flower colour and shape between *I. haynei* and *I. lortetii* (Sapir et al. 2002), these taxa are genetically close, and several *I. haynei* populations (s.s.) clustered with *I. lortetii* (Figure 3). Likewise, despite being morphologically well discriminated species (Danin and Fragman-Sapir 2016+), *I. petrana* and *atrofusca* were not assigned as separated genetic groups. These results are in accordance with a recent study, on a subset of the Israeli *Iris* species, which also found that *I. petrana* and *I. atrofusca* share the same gene pool (Volis et al. 2021).

These findings highlight the limitations of morphology-based taxonomy in recently diverged groups and emphasize the need for integrative approaches that combine genomic, ecological, and spatial data to accurately define species boundaries.

### The role of introgression and incomplete lineage sorting in shaping divergence

Differentiating between incomplete lineage sorting (ILS) and introgression is a central challenge in speciation genomics. particularly in young radiations where both processes can produce similar patterns of shared ancestry and phylogenetic discordance. In *Oncocyclus*, several species exhibit weak phylogenetic separation, non-monophyletic relationships, and overlapping genomic ancestry. Notably, *I. haynei*, *I. petrana*, and *I. atrofusca* form a phylogenetically intermixed group and share STRUCTURE components. Similarly, *I. mariae* and *I. atropurpurea* show substantial genetic overlap in STRUCTURE, despite forming distinct clades in the phylogeny. These patterns may result from either ILS or historical introgression.

One particularly interesting case involves the *I. hermona* SMK population, which clustered unexpectedly with *I. bismarckiana* in the phylogeny. This could reflect misidentification, ancestral polymorphism, or historical introgression. While ABBA-BABA tests found no significant introgression between SMK and *I. bismarckiana*, it revealed gene flow between *I. bismarckiana* and other *I. hermona* populations, suggesting past introgression. These findings support *I. bismarckiana* as a deeply divergent lineage, while also indicating possible historical gene flow with *I. hermona*. This interpretation is further supported by previous morphological work, which identify *I. bismarckiana* as a sister taxon to *I. hermona* (Sapir et al. 2001).

An unverified hypothesis suggests that *I. bismarckiana* was introduced to the region from the Damascus basin (Syria) by Crusaders in the 11th century (D. Shahak, personal communication). This was based on the species’ proximity to Crusader fortresses and its morphological resemblance to *I. damascene* (D. Shahak, pers. comm.). Testing such a hypothesis would require coalescent analyses and broader sampling of *I. bismarckiana*, *I. damascene*, and related taxa across the region.

Our ABBA-BABA results provide further insights into the historical dynamics of gene flow in *Oncocyclus*. In several cases of non-monophyly, such as among *I. haynei* and *I. atrofusca*, no significant introgression was detected, suggesting that ILS is the more likely explanation for their genetic overlap. In contrast, we found strong signals of introgression involving *I. mariae* and several other taxa. Significant excess allele sharing was detected between *I. mariae* and *I. atrofusca*, and even more so when *I. petrana* and southern *I. haynei* populations were included. These patterns suggest historical gene flow, potentially during periods of secondary contact following range expansion. Despite shared ancestry in STRUCTURE analysis, *I. atropurpurea* forms a well-supported monophyletic clade and shows no evidence of recent introgression.

These results also help clarify previous findings of admixture between *I. mariae* and *I. atropurpurea* and their relationships with *I. petrana* and *I. atrofusca*. Volis et al. showed some separation between *I. mariae* and the other species, but in accordance with our study, found shared genetic components between *I. mariae* and *I. atropurpurea*, and speculated that *I. atropurpurea* is of hybrid origin (Volis et al. 2021). However, our results show that *I. atropurpurea* is monophyletic and likely represents the ancestral lineage from which *I. mariae* diverged following southward dispersal. This interpretation is supported by strong ecogeographical reproductive isolation between *I. atropurpurea* and *I. mariae* (RI=0.99), and low isolation between *I. mariae* and *I. petrana* (RI=0.6-0.75), maintained by both ecological and phenological barriers (Osmolovsky et al. 2022). Overall, our findings support the presence of partial pre-zygotic isolation among these species, reported in previous ecological studies.

Taken together, these results highlight the complexity of divergence in *Oncocyclus*, where both ILS and historical gene flow can lead to unexpected phylogenetic patterns. This emphasizes the importance of integrating genomic, ecological, and spatial data to resolve evolutionary histories in young, rapidly diversifying species. To better understand the ecological and spatial context in which these divergence and introgression patterns occur, we examined how geography and environment structure genetic variation across the clade.

### The interplay of geography and ecology in driving divergence

Our results reveal that both geographic distance and environmental variation contribute to genetic divergence among *Oncocyclus* irises, consistent with patterns of isolation by distance (IBD) and isolation by environment (IBE). IBD remained significant across all taxa, while IBE became stronger once *I. bismarckiana* was excluded, with altitude, temperature, and aridity emerging as significant predictors of genetic structure. This suggests that both IBD and, to a lesser extent, IBE contribute to the speciation process in the Royal Irises, and that local adaptation may have reinforced divergence following initial spatial isolation. Our results support a model of speciation in which population dispersion and non-adaptive “stepping-stone” differentiation are followed by local adaptation to environmental gradients, potentially leading to ecogeographical reproductive barriers (Osmolovsky et al. 2022). This interpretation is consistent with a field study showing that divergence among *I. atropurpurea* populations is primarily driven by IBE, likely occurring after their dispersal (Yardeni et al. 2016). Additional support comes from common-garden studies showing that Royal Iris species exhibit local adaptation to soil and water availability, with species-specific differences in plasticity and performance on native versus non-native substrates (Dorman et al. 2009, Volis et al. 2021).

Together, these findings suggest that ecological divergence may act sequentially with spatial isolation to reduce gene flow and maintain species boundaries, even in the face of introgression. our results suggest that despite gene flow among certain *Oncocyclus* taxa, genomic regions associated with ecological or phenotypic divergence may be maintained by selection. The presence of well-defined clades alongside admixed lineages supports a scenario in which selection against maladaptive hybrids and local ecological differentiation jointly constrain introgression, preserving species identity.

### Implications for understanding speciation in complex landscapes

Taken together, our genomic analyses, spatial data, and ecological associations suggest a scenario of diversification in the *Oncocyclus* irises shaped by historical range expansion, ecological divergence, and localized gene flow (Figure 7). We found compelling evidence that range expansion of these plants from their putative origin in the North-East (Damascus basin and Caucasus) (Wilson et al. 2016) was accompanied by genetic differentiation and local adaptation to environmental factors such as climate and water availability. Our phylogenetic results suggest that three populations/species groups of diverged in parallel across the region (Figure 7). The first group includes the *I. atropurpurea-mariae* clade, in which *I. atropurpurea* spread along the coastal dunes and some individuals potentially dispersed southward to the arid desert sand dunes. This shift may have led to the emergence of, *I. mariae*, which exhibits low morphological variation (Sapir et al. 2002), suggesting a strong population bottleneck during adaptation to aridity. The second group includes *I. atrofusca*, *I. petrana*, and *I. haynei* (excluding the Gilboa populations, s.s.). We propose that *I. atrofusca* spread southward and speciated into *I. petrana*, and that these species came into (secondary?) contact with either *I. mariae*, or the southern populations of *I. atropurpurea*, leading to introgression that may have contributed to the distinct genomic profile of *I. mariae*. A third group of divergence includes the *I. haynei* s.s. (Gilboa populations) and *I. lortetii* clade. However, the spatial and temporal process of divergence in this clade remain unclear.

**Figure 7.**
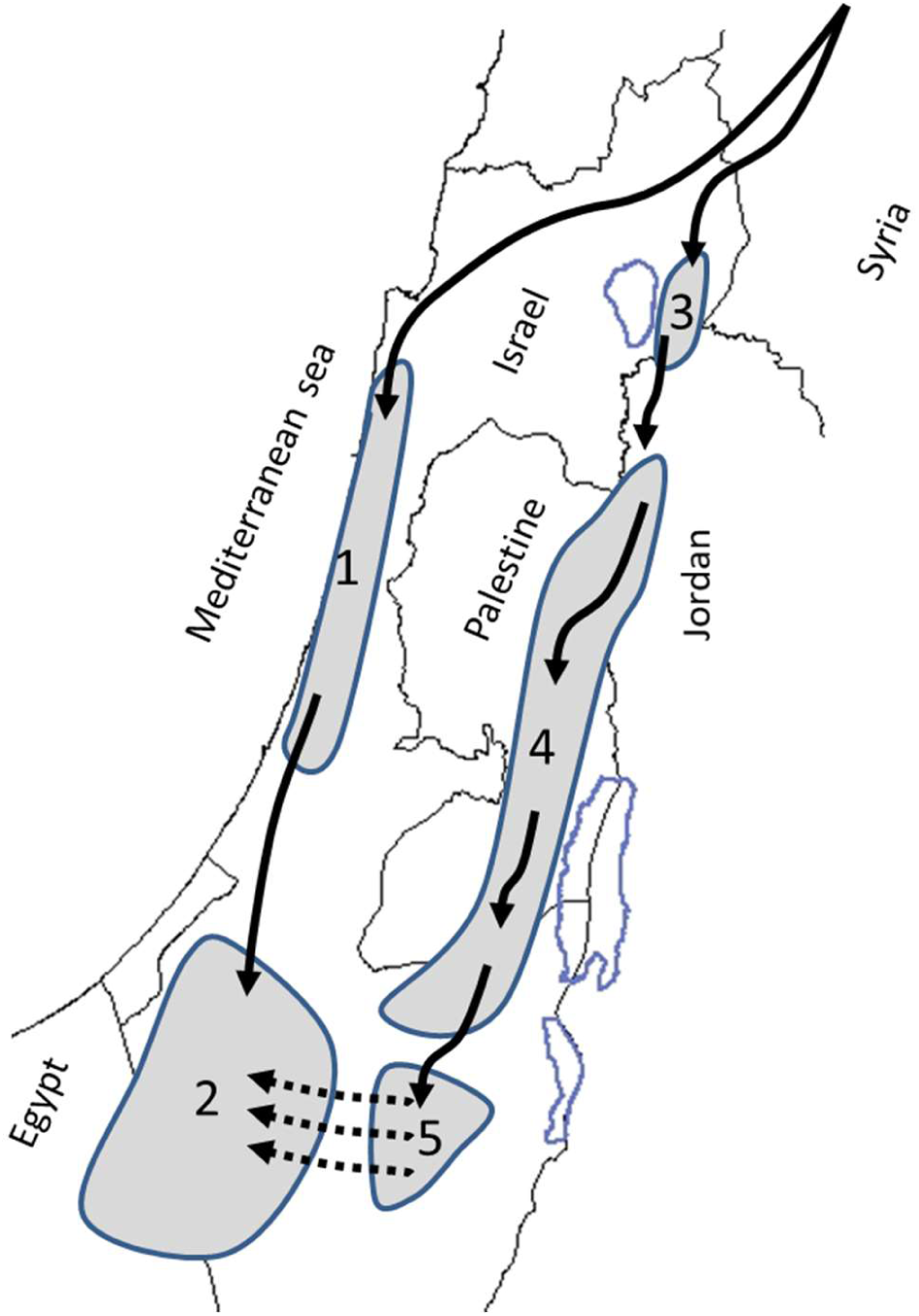
Hypothetical divergence process of *Oncocyclus* lineages in the southern Levant. Solid arrows – dispersion steps, possibly involving local adaptation. Dashed arrows – gene flow and introgression. 1 – *I. atropurpurea*; 2 – *I. mariae*; 3 – *I. haynei* sensu lato; 4 – *I. atrofusca*; 5 – *I. petrana*. Grey areas are the current distribution (see map in Figure 1). Species names are following the current taxonomy.

Similar patterns have been documented in other plant systems, where secondary contact following range expansion has led to localized introgression while preserving species boundaries. In *Senecio* species along an elevational gradient on Mount Etna, divergent QTL clusters and ecological selection maintained phenotypic and genomic divergence even in the face of gene flow (Brennan et al. 2016). In some species, such as *Senecio lautus*, rapid shifts to novel habitats have produced population bottlenecks and strong phenotypic differentiation (Roda et al. 2013). In the Louisiana irises, introgressed alleles improved hybrid fitness in specific environments, facilitating local adaptation while reproductive isolation was maintained through flowering time and ecological preferences (Bouck et al. 2005, Martin et al. 2005, 2006). Likewise, in *Helianthus*, hybridization among species after range expansion, has contributed to ecological divergence and the formation of hybrid species adapted to novel environments (Rieseberg et al. 1999). This is consistent with broader patterns in speciation genomics, where gene flow does not necessarily prevent local adaptation but may instead interact with selection in shaping heterogeneous genomic divergence (Tigano and Friesen 2016, Bock et al. 2023). These examples support our hypothesis that *Oncocyclus* irises underwent parallel divergence during range expansion, followed by secondary contact and limited introgression in ecologically distinct regions.

### Concluding Paragraph: Toward a predictive framework for speciation

Our results reveal that speciation in the *Oncocyclus* irises is shaped by a dynamic interplay of geographic isolation, ecological divergence, and historical gene flow, producing a complex pattern of genomic divergence. This study highlights the need for a multifactorial approach to speciation research, integrating genomic, ecological, and spatial data. Our findings contribute to the view of speciation as a heterogeneous and dynamic process, influenced by both barriers and bridges to gene flow (Abbott et al. 2013, Harrison and Larson 2014). Although focused on *Oncocyclus*, our findings reflect broader processes shaping speciation in many plant lineages, particularly those inhabiting topographically and environmentally heterogeneous regions. Future studies linking genomic divergence with phenotypic trait and reproductive barriers will be essential to understanding how speciation originates and proceeds in complex environments.

## Supporting information

BarLev_Supplementary material

## Acknowledgements

We thank I. Osmolovsky and K. Tielbürger for assistance in collecting materials and many discussions. We thank Dr. Eric Imbert from University of Montpellier, France, for providing plant material. This research was funded by the American Iris Society Foundation. S. Lozada-Gobilard was supported by a post-doctoral scholarship by University of Potsdam – Tel Aviv University collaboration program.

## Data accessibility

Raw RAD-Seq genetic data generated in this study have been deposited in the NCBI Sequence Read Archive (SRA), BioProject PRJNA995388. The VCF file containing the final SNP data can be found in the Supplementary.

## Benefit-Sharing Statement

All samples were collected with permission from Israel Nature Parks Authority to YS.

## Authors’ contributions

YBL and YS designed the study, collected the samples, and wrote the manuscript. YBL conducted the experimental work. YBL, SLG, AX, LG, and IM analyzed the data. All authors read and approved the final manuscript.

