## Supplementary material for "Evolutionary history, ecological divergence, and introgression in the *Oncocyclus* irises species complex in the Southern Levant": BarLev_Supplementary material

### TABLES

**Table S1.** Characteristics of the sampled populations.

### FIGURES

**Figure S1.** Genetic structure of *Iris* species

**Figure S2.** Discriminant Analysis of Principal Components (DAPC) of *Iris* species

**Figure S3.** Scatterplot of matrices of the environmental variables

**Figure S4.** Isolation by distance, Isolation by environment and RDA analysis for all species

**Table S1. Characteristics of the sampled populations.** Variables corresponding to Longitude and latitude were used to build a geographical matrix to test Isolation-by-Distance (IBD), while the environmental variables were used to build an ecological matrix to test Isolation-by-Environment (IBE). Long=Longitude, Lat=Latitude, MATemp=Mean Annual Temperature, MARain= Mean Annual precipitation, AEvap=Annual Evaporation, FebMinTemp=Minimal Temperature in February. Asterisks represent samples and localities that were excluded from ecological analyses due to missing ecological data.

| Species | Site | Location | RAD clusters | Long | Lat | Altitude | MATemp | MARain | AEvap | Aridity | FebMin Temp | Soil |
|---|---|---|---|---|---|---|---|---|---|---|---|---|
| *Iris atrofusca* Baker | ADD | Eldad | 92,605 | 35.248 | 31.656 | 598.28 | 18.81 | 303.21 | 1702.75 | 16.32 | 6.92 | Chalk |
| *Iris atrofusca* Baker | AND | Negev monument | 63,247 | 34.82 | 31.267 | 355.66 | 20.89 | 206.35 | 1592.92 | 9.86 | 8.76 | Loess |
| *Iris atrofusca* Baker | GRL | Goral | 142,347 | 34.798 | 31.31 | 327.89 | 20.47 | 223.63 | 1573.7 | 10.9 | 8.3 | Loess |
| *Iris atrofusca* Baker | MKH | Makukh | 42,310 | 35.36 | 31.905 | 307.43 | 21.52 | 216.4 | 1715.23 | 10.13 | 9.75 | Chalk, chert |
| *Iris atrofusca* Baker | MKS | Maale Mikhmas | 53,424 | 35.31 | 31.881 | 579.11 | 18.87 | 309.76 | 1675.52 | 16.81 | 6.68 | Chalk |
| *Iris atrofusca* Baker | MRT | Marit wadi | 126,195 | 35.03 | 31.258 | 440.17 | 19.56 | 190.49 | 1720.94 | 9.69 | 7.03 | Loess |
| *Iris atrofusca* Baker | RMN | Rimonim | 84,011 | 35.332 | 31.915 | 555.9 | 19.11 | 275.34 | 1681.68 | 14.34 | 6.75 | Chalk |
| *Iris atrofusca* Baker | TKO | Tekoa | 113,294 | 35.236 | 31.645 | 558.84 | 19.1 | 318.2 | 1698.64 | 16.94 | 7.3 | Limestone, dolomite, chert |
| *Iris atrofusca* Baker | TLA | Tel Arad | 120,939 | 35.122 | 31.273 | 522.31 | 18.97 | 167.43 | 1768.35 | 9.01 | 6.34 | Loess |
| *Iris atrofusca* Baker | YAR | Mizpe Yair | 41,976 | 35.146 | 31.381 | 743.13 | 18.4 | 224.15 | 1750.92 | 12.18 | 6.83 | Limestone |
| *Iris atropurpurea* Disnm. | ASH | Ashdod | 43,784 | 34.698 | 31.833 | 38.85 | 19.71 | 512.75 | 1427.54 | 25.94 | 8.2 | Sand |
| *Iris atropurpurea* Disnm. | DR | Dora, Netanya | 85,273 | 34.847 | 32.288 | 35.66 | 20.2 | 526.46 | 1407.91 | 26.04 | 9.73 | Sand |
| *Iris atropurpurea* Disnm. | HUM | Humra | 37,518 | 34.744 | 31.935 | 15.15 | 19.35 | 493.28 | 1426.66 | 25.31 | 7.26 | Hamra |
| *Iris atropurpurea* Disnm. | IAR | I.atropurpurea reserve | 106,736 | 34.749 | 31.93 | 31.22 | 19.46 | 497.28 | 1427.71 | 25.55 | 7.25 | Sand |
| *Iris atropurpurea* Disnm. | KDM | Kadima | 83,068 | 34.921 | 32.287 | 64.06 | 19.74 | 609.34 | 1382.61 | 31.05 | 7.96 | Sand |
| *Iris atropurpurea* Disnm. | KUR | Kur | 70,086 | 34.709 | 31.891 | 12.58 | 19.04 | 481.39 | 1424.57 | 25.06 | 7 | Sand |
| *Iris atropurpurea* Disnm. | NET | Netanya | 76,193 | 34.841 | 32.286 | 28.45 | 20.39 | 519.91 | 1407.73 | 25.51 | 10.48 | Sand |
| *Iris atropurpurea* Disnm. | NRG | Nir Galim | 84,481 | 34.674 | 31.829 | 13.24 | 19.87 | 496.22 | 1424.5 | 25.02 | 8.26 | Sand |
| *Iris atropurpurea* Disnm. | NSZ | Nes Ziona | 78,610 | 34.784 | 31.937 | 63.91 | 20.07 | 524 | 1432.04 | 26.04 | 8.47 | Sandstone |
| *Iris atropurpurea* Disnm. | NTA | Netaim | 103,600 | 34.762 | 31.946 | 31.85 | 19.33 | 504.61 | 1428.98 | 25.61 | 7.33 | Hamra |

| | | | | | | | | | | | | |
|---|---|---|---|---|---|---|---|---|---|---|---|---|
| *Iris atropurpurea* Disnm. | OLG | Olga | 67,089 | 34.877 | 32.433 | 11.96 | 19.48 | 502.54 | 1414.5 | 25.86 | 8.46 | Sand |
| *Iris atropurpurea* Disnm. | PLG | Poleg | 92,805 | 34.838 | 32.264 | 30.02 | 19.96 | 522.72 | 1407.58 | 26.01 | 9.17 | Sandstone |
| *Iris atropurpurea* Disnm. | RLZ | Rishon LeZion | 149,916 | 34.797 | 31.943 | 54.73 | 20.32 | 535.16 | 1433.27 | 26.4 | 8.58 | Hamra |
| *Iris atropurpurea* Disnm. | SHF | Shafdan | 73,116 | 34.751 | 31.96 | 25.97 | 19.6 | 491.98 | 1425.96 | 25.59 | 8.19 | Sand |
| *Iris atropurpurea* Disnm. | TLV | Tel Aviv | 60,191 | 34.809 | 32.133 | 20 | 20.07 | 536.83 | 1407.38 | 26.77 | 8.66 | Sandstone |
| *Iris atropurpurea* Disnm. | YQM | Yaqum | 62,529 | 34.844 | 32.254 | 19.84 | 19.62 | 531.15 | 1407.97 | 27.02 | 8.27 | Sandstone |
| *Iris auranitica* Disnm.* | JDZ | Jabel Druz | 82,551 | 36.715 | 32.715 | NA | NA | NA | NA | NA | NA | Basalt |
| *Iris bismarckiana* Regel | DSH | Dishon | 78,914 | 35.509 | 33.078 | 460.53 | 18.86 | 600.35 | 1471.88 | 32.57 | 6.85 | Chalk, marl |
| *Iris bismarckiana* Regel | GIV | Givat Hamore | 29,816 | 35.358 | 32.616 | 469.64 | 19.94 | 468.15 | 1560.38 | 23.19 | 8.85 | Limestone, chert |
| *Iris bismarckiana* Regel | HYN | Har Yona | 37,072 | 35.34 | 32.724 | 545.8 | 19 | 600 | 1522.42 | 31.5 | 7.73 | Limestone, chert |
| *Iris bismarckiana* Regel | MHZ | Mahoz (Tefen) | 45,935 | 35.271 | 32.949 | 445.23 | 18.71 | 844.91 | 1474.92 | 45.14 | 7.11 | Limestone |
| *Iris bismarckiana* Regel | MSG | Misgav wadi | 49,810 | 35.566 | 33.255 | 449.94 | 18.93 | 803.89 | 1441.47 | 43.04 | 7.17 | Limestone, chert |
| *Iris bismarckiana* Regel | NZR | Nazarat | 58,885 | 35.339 | 32.71 | 448.85 | 19.71 | 595.35 | 1522.32 | 30.44 | 8.12 | Chalk |
| *Iris bismarckiana* Regel * | RJW | Rajib | 54,165 | 35.693 | 32.401 | NA | NA | NA | NA | NA | NA | Chalk and marl |
| *Iris bismarckiana* Regel * | SMS | Majdal Shams | 73,695 | 35.774 | 33.278 | 1272.59 | 14.57 | 1199.83 | NA | 85.86 | 2.94 | Limestone, chert |
| *Iris bismarckiana* Regel | YFT | Yiftach | 27,130 | 35.551 | 33.139 | 434.83 | 18.53 | 579.33 | 1465.19 | 31 | 6.21 | Limestone, dolomite |
| *Iris haynei* Baker | BRK | Barkan | 86,097 | 35.411 | 32.506 | 450.38 | 19.7 | 393.2 | 1650.8 | 19.9 | 7.95 | Limestone, chert |
| *Iris haynei* Baker | HMD | Hemdat | 38,329 | 35.527 | 32.251 | 153.05 | 22.16 | 233.41 | 1714.94 | 10.58 | 9.84 | Limestone, chert |
| *Iris haynei* Baker | MGL | Maale Gilboa | 114,943 | 35.411 | 32.477 | 419.99 | 19.37 | 391.18 | 1664.09 | 20.36 | 7.16 | Limestone, chert |
| *Iris haynei* Baker | MVH | Mevo-Hamma, lower pop | 65,074 | 35.652 | 32.705 | 310.14 | 21.04 | 406.21 | 1609.05 | 19.15 | 8.89 | Basalt |
| *Iris haynei* Baker | PLS | Peles | 90,074 | 35.519 | 32.297 | 89.11 | 22.49 | 242.25 | 1706.79 | 10.78 | 10.18 | Sand |
| *Iris haynei* Baker | SGL | Mevo-Hamma, cliffs | 112,392 | 35.659 | 32.749 | 310.59 | 20.32 | 423.24 | 1601.38 | 20.26 | 8.45 | Basalt |
| *Iris haynei* Baker | SUA | Malkishua | 65,977 | 35.413 | 32.435 | 467.45 | 19.37 | 400.61 | 1665.28 | 20.62 | 7.77 | Limestone, chert |
| *Iris haynei* Baker | ZRR | Mezar | 51,797 | 35.683 | 32.767 | 287.37 | 20.61 | 460.73 | 1604.6 | 22.21 | 8.18 | Basalt |
| *Iris hermona* Disnm. | KST | Keshet | 125,940 | 35.805 | 32.981 | 697.67 | 16.76 | 673.26 | 1529.29 | 40.23 | 4.47 | Basalt |
| *Iris hermona* Disnm. | MPL | Mapalim | 79,158 | 35.752 | 32.987 | 531.02 | 18.43 | 611.61 | 1528.85 | 32.95 | 5.93 | Basalt |
| *Iris hermona* Disnm. | MRZ | Shekh Marzuk | 83,736 | 35.703 | 33.049 | 526.09 | 18.06 | 602.83 | 1500.4 | 33.31 | 5.42 | Basalt |

| | | | | | | | | | | | | |
|---|---|---|---|---|---|---|---|---|---|---|---|---|
| *Iris hermona* Disnm. | SMK | Samak gorge, Golan | 152,544 | 35.715 | 32.833 | 71.71 | 21.84 | 505.2 | 1590.71 | 23.03 | 9.25 | Chalk |
| *Iris lortetii* Barbey * | AVV | Avivim | 88,549 | 35.501 | 33.088 | 683.77 | 17.44 | 620.68 | NA | 36.37 | 5.76 | Limestone, chert |
| *Iris lortetii* Barbey | AYL | Ayelet Hashahar | 57,106 | 35.567 | 33.014 | 229.49 | 20.23 | 510.89 | 1511.16 | 25.36 | 7 | Sand |
| *Iris lortetii* Barbey | BTD | Beit Dajan | 24,669 | 35.4 | 32.19 | 637 | 18.68 | 375.86 | 1657.95 | 20.2 | 7.31 | Limestone, dolomite, chert |
| *Iris lortetii* Barbey | KDS | Nahal Kedesh | 39,522 | 35.553 | 33.117 | 364.58 | 19.21 | 524.25 | 1470.61 | 27.51 | 6.94 | Limestone |
| *Iris lortetii* Barbey | MLK | Malkia | 96,492 | 35.518 | 33.11 | 581.65 | 17.8 | 589.25 | 1464.4 | 33.18 | 5.69 | Limestone, chert |
| *Iris lortetii* Barbey | MNR | Manara | 50,090 | 35.557 | 33.199 | 430.36 | 19.1 | 692.82 | 1451.84 | 37.31 | 7.92 | Limestone |
| *Iris lortetii* Barbey | PUA | Pua Mt. | 52,878 | 35.451 | 33.052 | 618.6 | 17.33 | 733.27 | 1462.28 | 41.91 | 5.68 | Limestone, chert |
| *Iris mariae* Barbey | BER | Beer Malca | 123,364 | 34.412 | 30.938 | 225.08 | 20.6 | 75.02 | 1573.2 | 3.64 | 8.27 | Sand |
| *Iris mariae* Barbey | HAL | Haluzit | 30,254 | 34.343 | 31.171 | 102.98 | 20.13 | 147.88 | 1518.64 | 7.35 | 7.72 | Sand |
| *Iris mariae* Barbey | NYZ | Nir Ytzhak | 57,192 | 34.358 | 31.241 | 98.93 | 19.96 | 190.38 | 1504.5 | 9.53 | 7.74 | Sand |
| *Iris mariae* Barbey | SHV | Shivta | 30,706 | 34.606 | 30.954 | 318.23 | 19.22 | 91.8 | 1589.15 | 4.75 | 6.72 | Sand |
| *Iris mariae* Barbey | SKR | Nahal Sekher | 57,490 | 34.8 | 31.12 | 310.52 | 19.42 | 128.16 | 1631.64 | 6.47 | 6.77 | Chalk |
| *Iris mariae* Barbey | ZEL | Zeelim | 56,818 | 34.512 | 31.193 | 145.15 | 19.97 | 153.13 | 1525.38 | 7.67 | 7.37 | Sand |
| *Iris petrana* Disnm. | DMN | Dimona | 92,470 | 35 | 31.07 | 518.76 | 18.71 | 118.99 | 1780.41 | 6.33 | 6.22 | Sand |
| *Iris petrana* Disnm. * | DNA | Jordan | 35,022 | 35.614 | 30.68 | NA | NA | NA | NA | NA | NA | Limestone |
| *Iris petrana* Disnm. | EFE | Efeh dunes | 32,606 | 35.145 | 31.063 | 331.29 | 20.7 | 73.18 | 1912.09 | 3.52 | 8.04 | Sand |
| *Iris petrana* Disnm. | MMS | Mamshit | 86,499 | 35.065 | 31.035 | 470.01 | 19.81 | 92.5 | 1848.45 | 4.7 | 7.62 | Sand |
| *Iris petrana* Disnm. | YER | Yeruham | 126,674 | 34.987 | 31.027 | 557.28 | 18.27 | 110.89 | 1788.83 | 6.07 | 5.82 | Sand, loess |
| *Iris petrana* Disnm. | YMN | Mishor Yamin | 63,096 | 35.126 | 31.039 | 398.28 | 20.17 | 72.79 | 1903.62 | 3.59 | 7.53 | Sand |
| *Iris petrana* Disnm. | ZFT | Zafit | 49,874 | 35.19 | 31.041 | 411.82 | 20.24 | 64.22 | 1963.76 | 3.16 | 7.6 | Sand |
| *Iris mesopotamica* Dykes | MES | Mount Hermon, Isarel | 76,850 | | | | | | | | | |
| *Iris lutescens* Lam | P1 | Gardiole, France | 105,404 | | | | | | | | | |
| *Iris lutescens* Lam | P4 | BelAir, France | 139,736 | | | | | | | | | |
| *Iris lutescens* Lam | Y5 | BelAir inconnue, France | 60,948 | | | | | | | | | |
| *Iris lutescens* Lam | Y6 | St Jean, France | 76,326 | | | | | | | | | |

### FIGURES

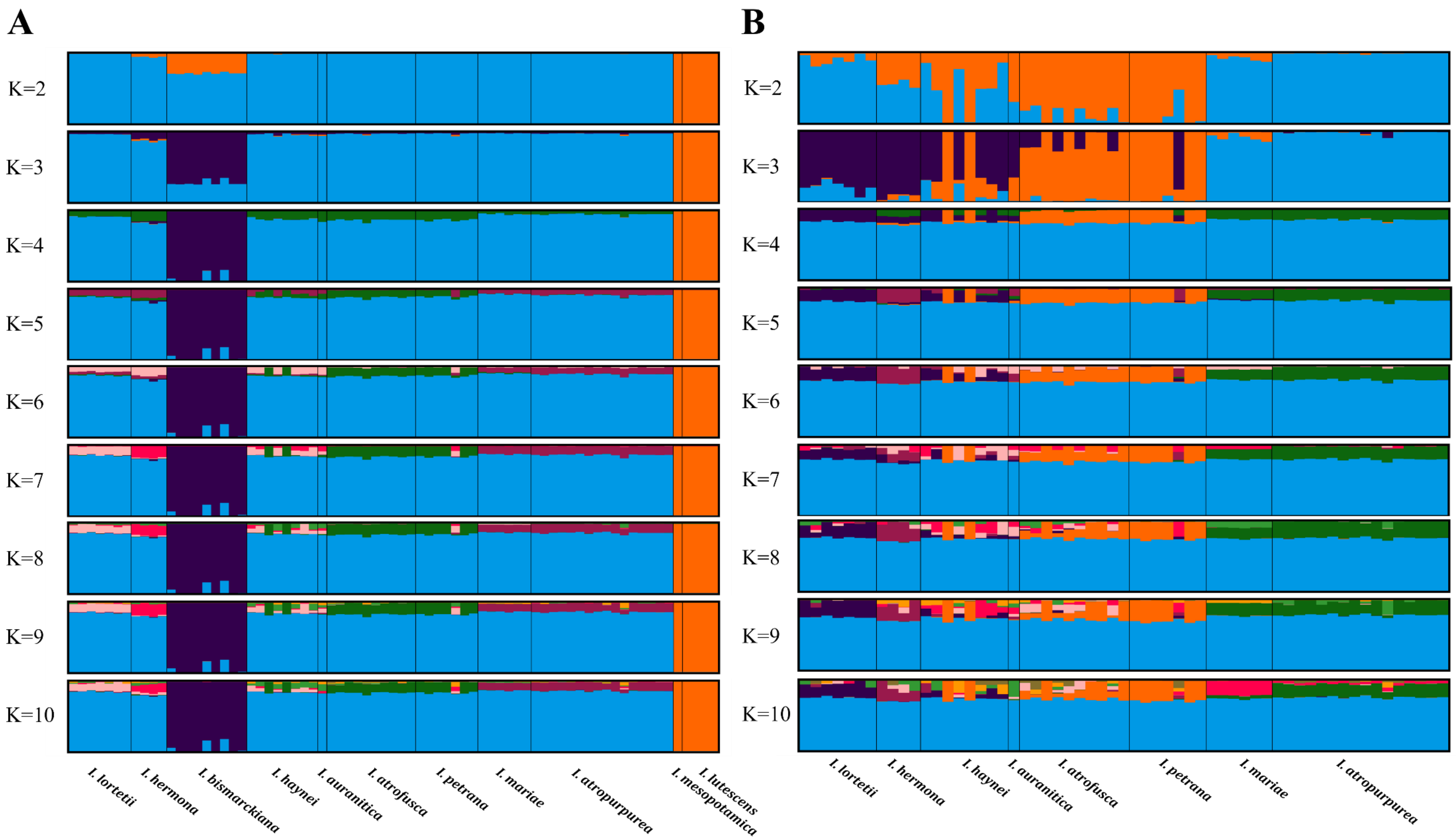


**Figure S1. Genetic structure of *Iris* species.** STRUCTURE plots of all populations (n=73) **(A)** and excluding the outgroups and *I. bismarckiana* (n=59) **(B)**. Ten independent runs were clustered and averaged using CLUMPAK for each *K* from 2 to 10. A single vertical line represents each individual; a black line separates species; the whole sample is divided into colors representing the number of clusters assumed.

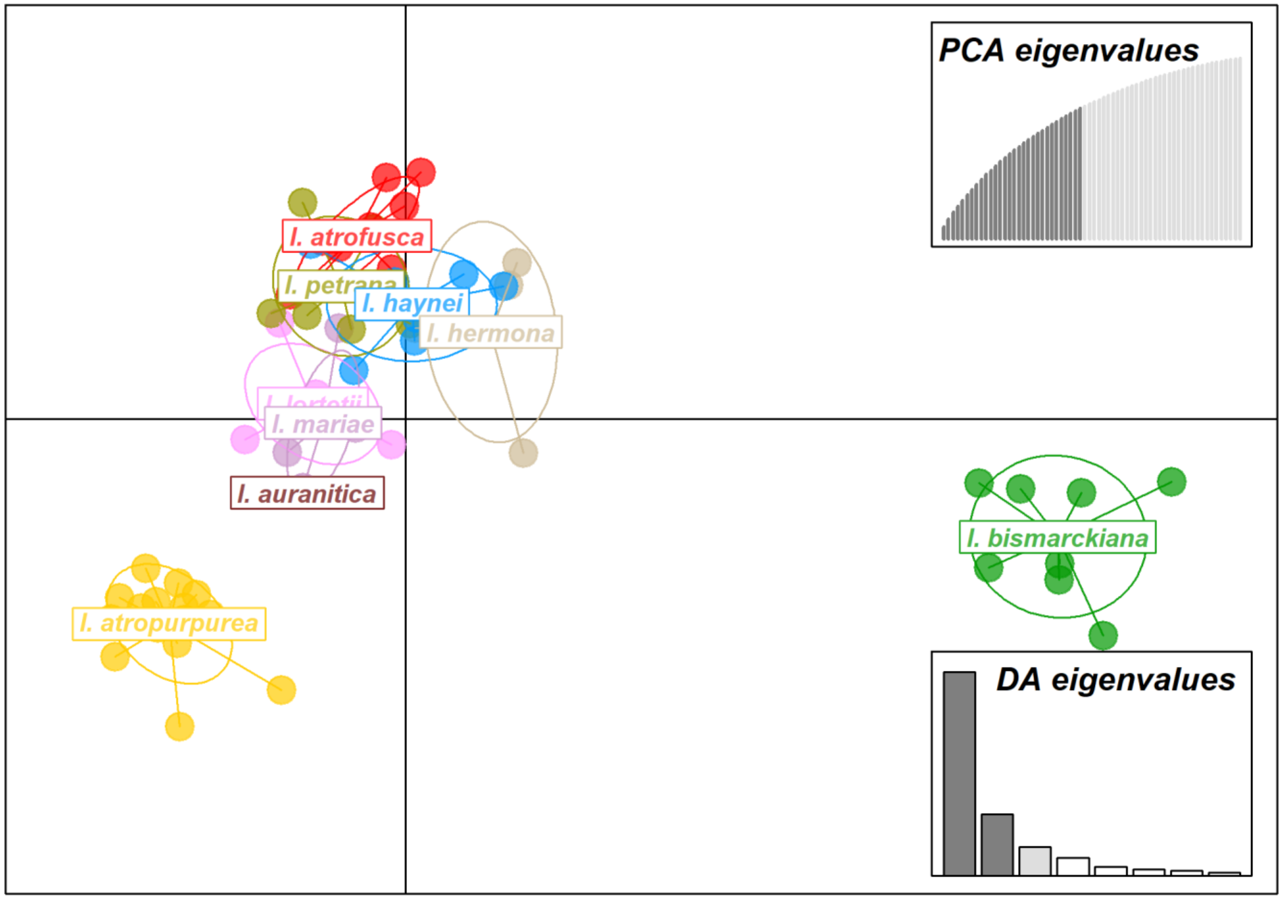


**Figure S2. Discriminant Analysis of Principal Components (DAPC) of *Iris* species**. DAPC plot including *I. bismarckiana* but excluding the outgroup species (*I. lutescens* and *I. mesopotamica)* (n=68). DAPC was based on 30 retained principal components (explaining 72.1% of total variance) and 3 discriminant functions (explaining 87.4% of between-group variation). *Iris bismarckiana* forms a highly distinct and isolated cluster, while the remaining taxa exhibit partial clustering and some overlap. This plot highlights the outlier position of *I. bismarckiana* and supports its exclusion in subsequent DAPC analyses to better resolve finer-scale structure.

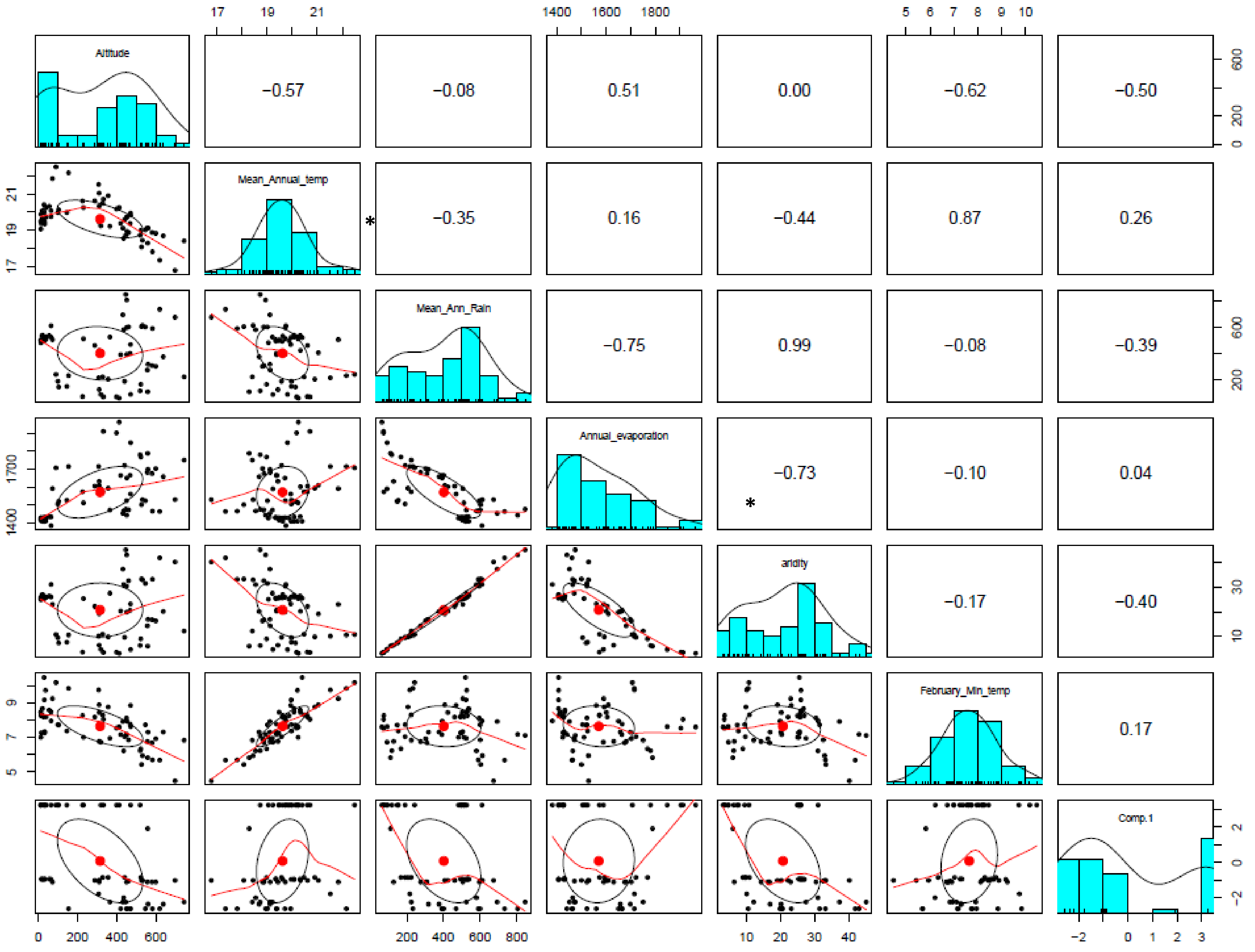


**Figure S3. Scatterplot of matrices of the environmental variables.** Bivariate scatterplots are shown below the diagonal, histograms on the diagonal, and the Pearson correlation above the diagonal. Asterisks represent variables that were excluded from ecological analyses due to high correlation (|r|>0.75).

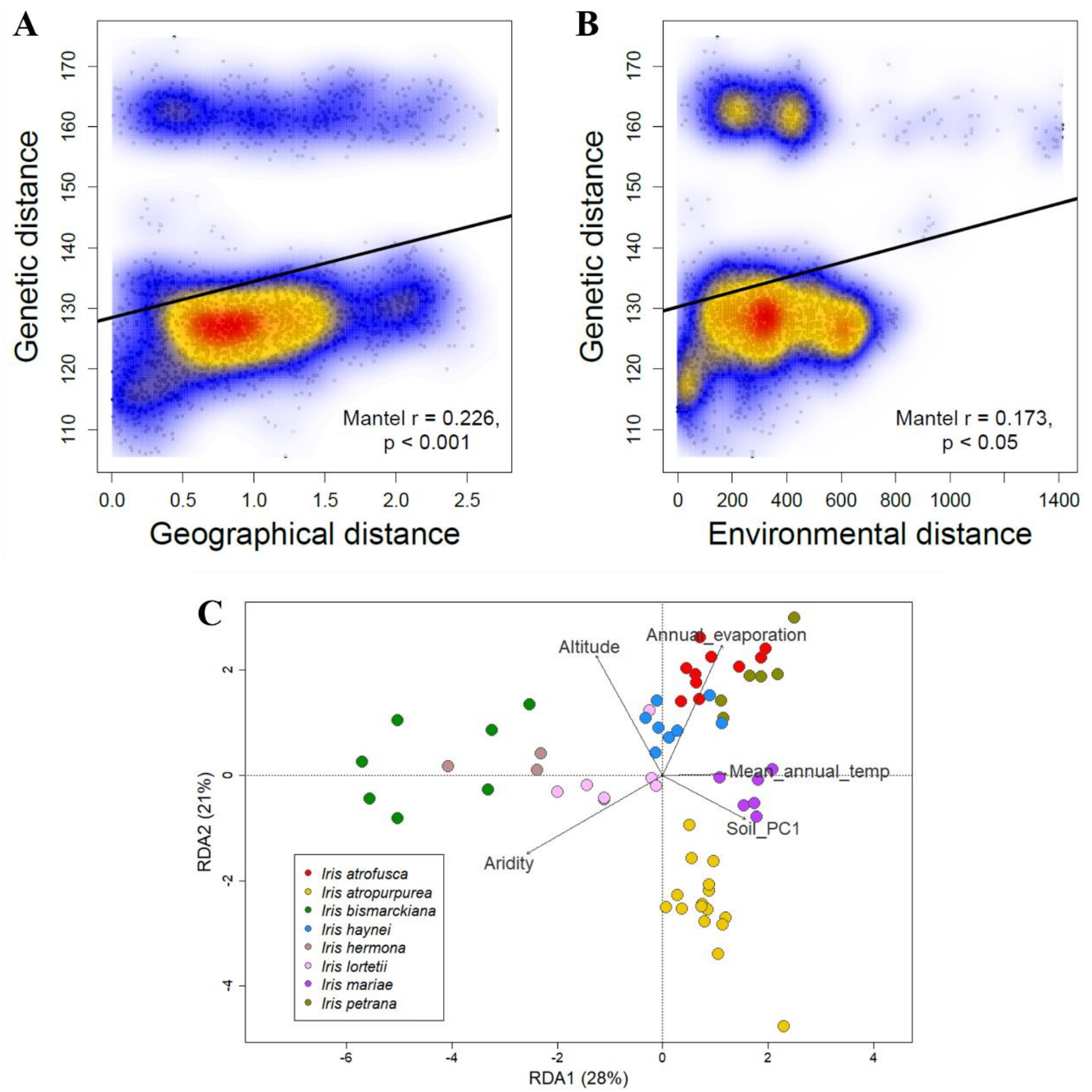


**Figure S4.** Isolation by distance **(A)** and Isolation by environment **(B)** calculated by Mantel test across all species (n=63). Each dot represents a pair of populations, with position indicating their genetic distance and environmental similarity. Colors denote density of points. The solid line within each panel is the linear regression line. Mantel coefficient (ρ) of correlation between the matrices, and the p-values are displayed at the bottom right hand side of each panel. **(C)** Redundancy analysis (RDA) ordination plot of the environmental variables based on genetic distance between all *Iris* species examined. Each point represents a population, colours indicate the species, and the arrows represent the environmental variable. For abbreviations meaning refer to Table S1.
